# Hybridization Reveals Cell Type-Specific Regulatory Variation Driving Brain Transcriptomic Divergence

**DOI:** 10.1101/2025.11.04.686555

**Authors:** Erika Soria, X Maggs, Will Boswell, Rachel A. Carroll, Korri S Weldon, Zhao Lai, Wesley C. Warren, Manfred Schartl, Yuan Lu

**Author notes:** Corresponding authors: Yuan Lu.

## Abstract

Gene expression is a molecular trait that can cumulatively contribute to more complex phenotypes. Interspecies hybrids have long been used to study the genetics basis of molecular, morphological and behavioral traits, to dissect interactions between cis- and trans-regulators with parental traits, and to uncover incompatible genetic interactions that drive extreme phenotypes in hybrid progeny. However, it remains unclear how organ functions at molecular level are affected by hybridization. We hypothesize that hybridization incited molecular level changes are cell type specific. To test this, we produced interspecies hybrid progeny from two distantly related fish species, *Xiphophorus maculatus* and *couchianus*. They are differing in mating, foraging behavior and likely neural circuits for cognition. We first performed allelic expression and bulk brain transcriptome profiling to identify expression quantitative trait loci (eQTLs) and quantitative transcript traits (QTTs) in order to quantify the scale and identify of loci contributing to gene expression variations in hybrids. It showed that QTT are predominantly influenced by additive eQTLs. Subsequently, we compared QTTs, which exemplify species-specific regulatory effects on gene expression, to cell type markers derived from single-nucleus RNA sequencing (snRNAseq). We identified 14 cell type-specific QTTs with known roles in brain functions. Overall, this study shows that transcriptomic phenotypes under species-specific regulators are associated to specific cell types in hybrids, and indicates the overall organ-level functional change could be driven by particular cell types.

## Introduction

Gene expression is a molecular phenotype underlying complex morphological and behavioral traits. Genetics variants contribution to gene expression has been studied for more than two decades using expression quantitative trait locus (eQTL) studies (1). The proportion of the hybrid genome exhibiting regulated and dysregulated gene expression, and the contribution of cis- and trans-regulators to gene expression have been investigated in a few model organisms involving hybrids produced from distant populations, such as different species (2–12). Hybrid dysgenesis and hybrid vigor focus on measurable phenotypic traits, such as morphology, fecundity, reproductive fitness, and life history traits. However, changes in molecular traits, such as gene expression, underlying whole organ functional difference in interspecies hybrid is understudied.

It has been reported in human studies that eQTL exhibit their influence on gene expression in a cell type-specific manner (13–15). These observations lead to the hypothesis that eQTL and quantitative trait transcripts (QTT; genes under eQTL regulation) are cell type-specific, and that organ level function in hybrid is contributed by certain cell types within an organ. Specifically, genetics underpinnings on brain function, such as cognition, foraging and mating behavior is poorly studied. Therefore, in this study, we aim to investigate spatiality of parental-specific molecular traits by locating cell types exhibiting QTTs.

*Xiphophorus* fishes serve as a vertebrate system to study genetics underlying divergent phenotypes. There are adequate resources available to this model system, including domesticated animals, sequenced reference genomes, documented genetics, morphological and behavioral divergences, and capability to produce an interspecies hybrid test population (16). We used hybrids between two distantly related parental species showing diverged behavior traits, *X. maculatus* and *X. couchianus* (Supp. Fig. 1). We performed bulk brain RNAseq and parental snRNAseq that enabled cell context by genotype to study gene(s) individual contributions. Accordingly, we assessed the genomic scale and genetic identity of brain QTT and regulatory inter-specific genome variants as eQTLs, and subsequently deconvoluted these expression phenotypes into brain cell types.

## Results

### Genetic interaction in the *Xiphophorus* brain

A total of 109 F_2_ interspecies hybrids whole brain samples were sequenced for genotyping and bulk gene expression profiling. Total numbers of quantitative trait transcript (QTT) and expression quantitative trait loci (eQTL) exhibiting additive, and dominant are identified (Fig. 1; -log_10_ adj. p-val > 10).

**Figure 1.**
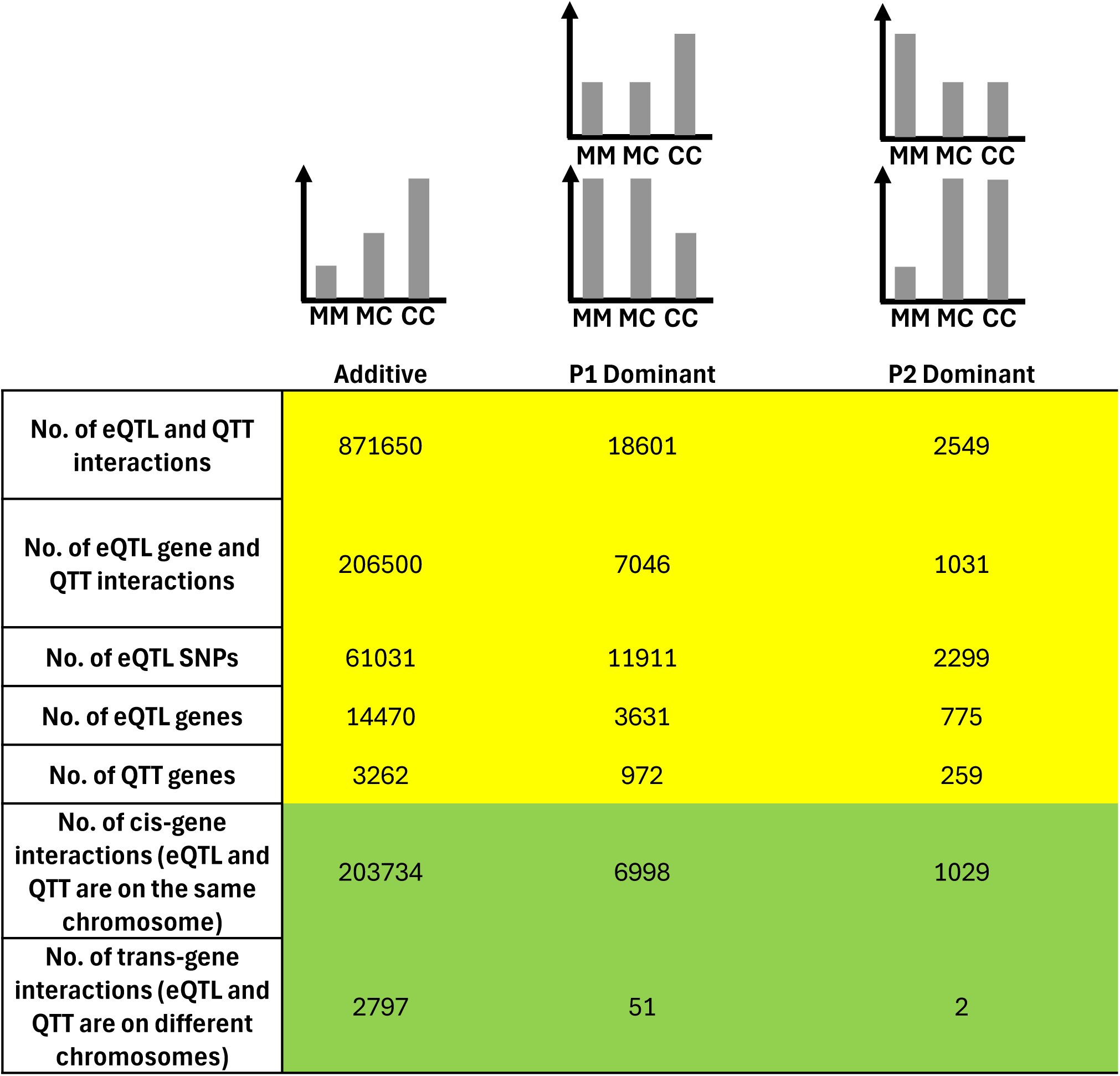
Quantification of eQTLs and QTTs. Three types of eQTLs were identified: additive, dominant for *X. maculatus* (P1) allele (M allele), and dominant for *X. couchianus* (P2) allele (C allele) using an FDR adjusted ANNOVA -log_10_p-value > 10. Yellow colored table reports quantity of eQTL-QTT interactions at inter-species polymorphism and gene levels. Green colored table reports quantity of cis- and trans-genetic interactions of each type.

Additive eQTLs dominate the dataset (97.6%; Fig. 1). For eQTLs that exhibited parental allele dominance, a majority of them (87.6%; Fig. 1) are from the *X. maculatus* parental allele.

Earlier studies comparing F_1_ hybrids to parentals showed there were globally distributed gene dysregulation (17). However, there is no over-dominance and very few under-dominance eQTLs, even with a lowered statistical stringency (-log_10_ adj. p-value > 2; Supp. Fig. 2). Mitonuclear genome incompatibility was reported to be lethal in certain *Xiphophorus* hybrids (18). All hybrids in this study inherited *X. maculatus* maternal mitochondria due to the hybridization scheme (Supp Data-MT genotype). Therefore, only nuclear polymorphic sites’ genotypes association with mitochondrial gene expression were assessed. Only under-dominant nuclear genome eQTLs associated with 5 mitochondrial QTTs were identified with relatively lowered statistical stringency (-log_10_ adj. p-val > 2; Supplement Fig. 3).

The genomic positions of the eQTL and QTTs showed that a substantial proportion of the genetic interactions are located on the same chromosome and are close to each other (Fig.1; Fig. 2). This is reflected in the typical eQTL-QTT distribution along the diagonal stripe of the scatter plot (Fig. 2). The eQTLs form clusters that are rather large. We measured an average additive eQTL cluster size of 8.1 Mbp (Supp. Fig 4) that is equivalent to 12 genes. Such eQTL cluster sizes were observed in other studies adopting inter-subspecies hybrids, such as hybrids between *M. musculus* subspecies (19). In comparison, a minor proportion of additive and dominant eQTLs locate on different chromosomes from QTTs (i.e., trans-interaction; Fig. 1).

**Figure 2.**
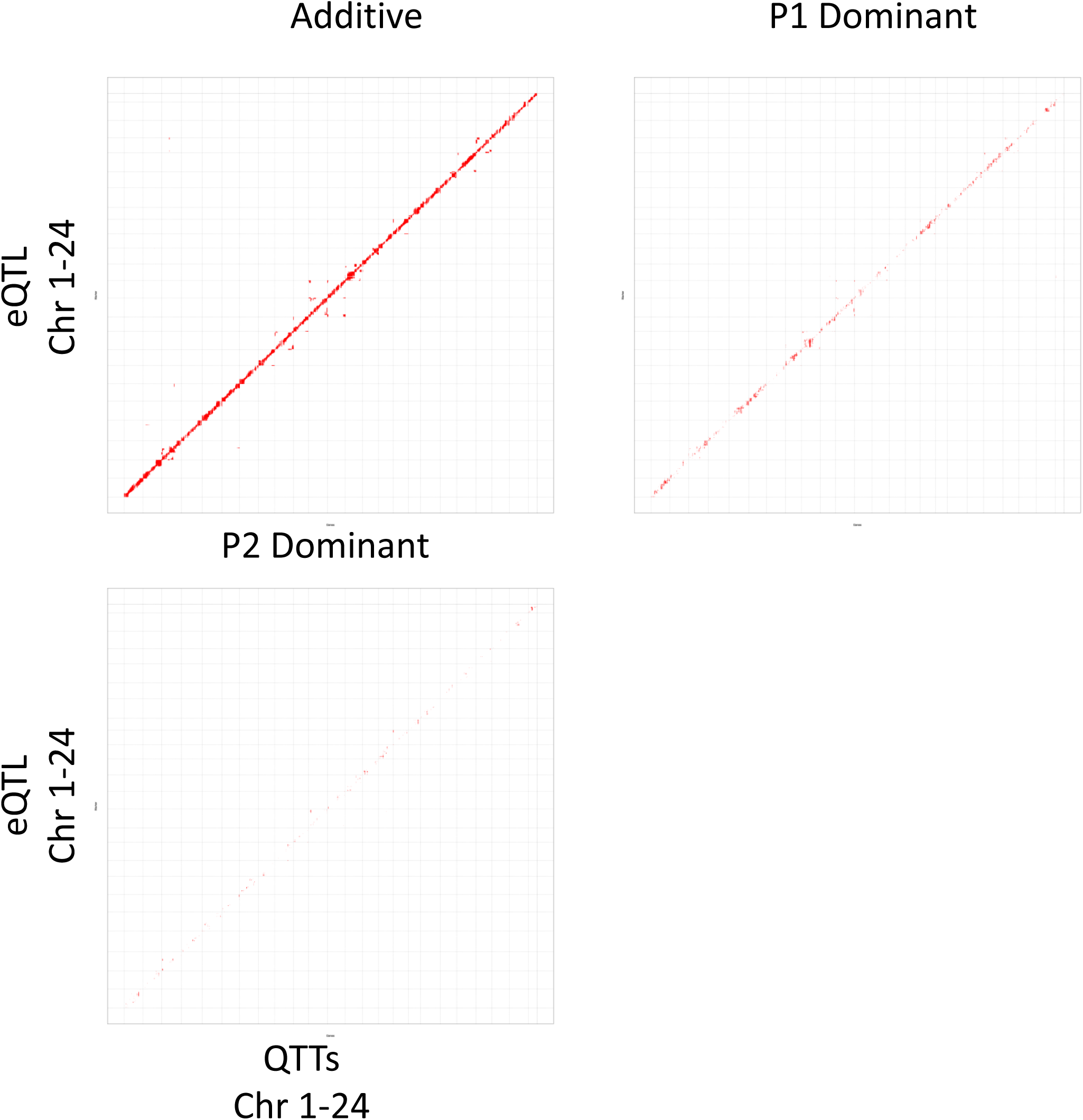
Genomic coordinates of eQTLs and QTTs. Scatter plots show genomic coordinates of additive and dominant eQTLs and QTTs. The plots are separated by vertical and horizontal lines, which hallmark the chromosome boundaries. Red dots represent statistically significant eQTL-QTT interactions. The X-coordinate of a red dot corresponds to physical location of an eQTL, and Y-coordinate corresponds to physical location of associated QTT.

### Deconvoluting bulk brain expression into cell types

To study cell type specific QTTs, snRNAseq was conducted on the parental *X. maculatus* brain. A total of 17,898 nuclei were sequenced, and 14 transcriptionally distinct clusters were generated. By comparing to expressed genes that define zebrafish and mouse brain cell types, 13 of the 14 clusters were annotated. The major cell types identified included neurons, astrocytes, oligodendrocytes, microglial cells and endothelial cells (Fig. 3; Supplement Table Cell type annotation).

**Figure 3.**
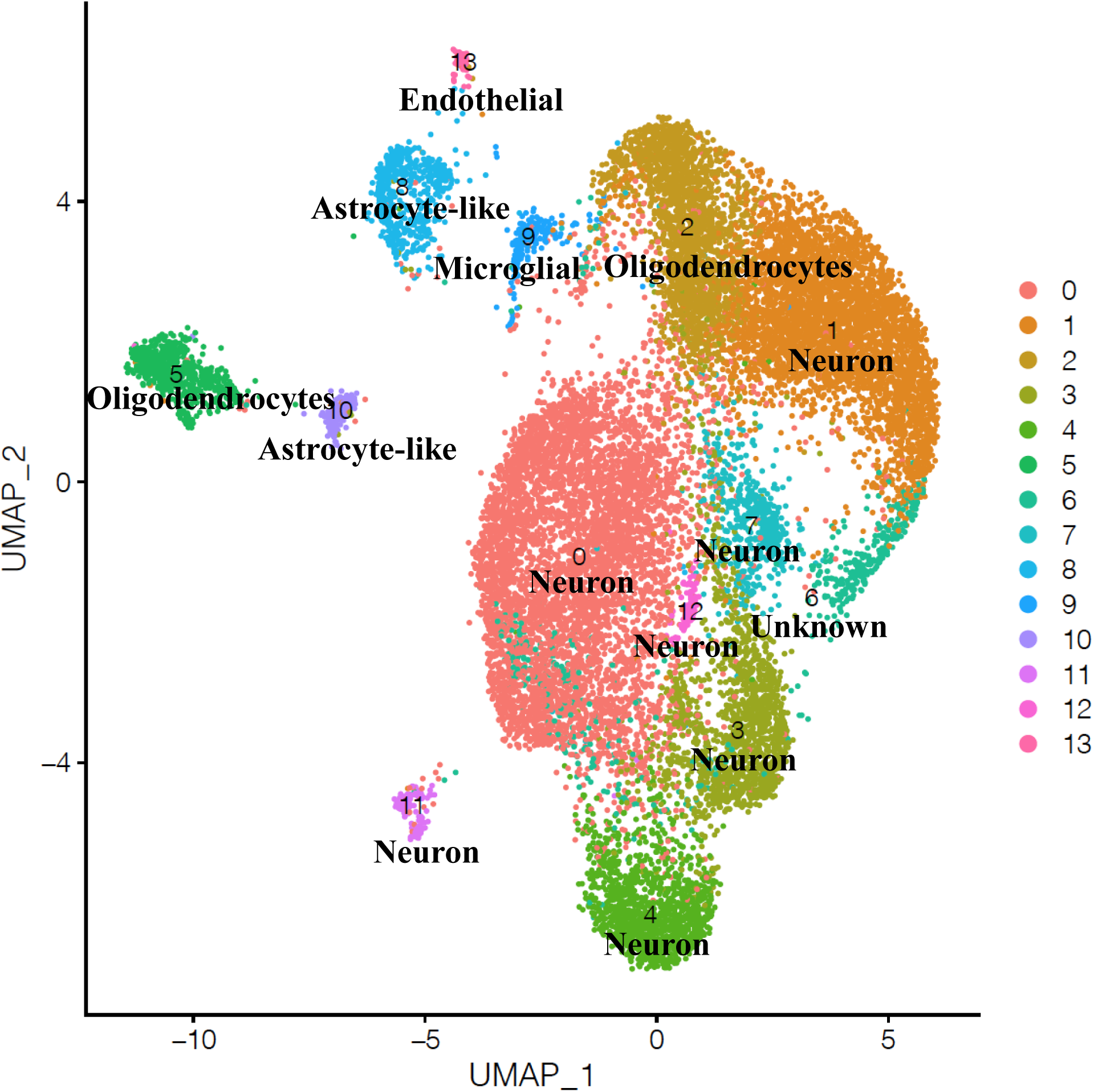
Cell types identified in *Xiphophorus* brain. The snRNAseq data, following gene expression quantification, is dimension reduced and plotted on Uniform Manifold Approximation and Projection (UMAP) for visualization. By comparing markers of each cell cluster to known brain cell type marker genes in zebrafish and mouse, 13 of the 14 cell clusters can be identified. Each dot on the UMAP represent a sequenced nucleus, with the color represented identity of cluster and cell type.

Nuclei expressing QTTs were identified and tested for marker gene enrichment by cell type (average log_2_FC >0, adjusted p-val <0.01, percentage of cell expressing a QTT within a cell type > 20% and a ratio between percentage of cell expressing a QTT within a cell type over percentage of cell expressing a QTT in the rest of the whole dataset > 5; Fig. 4; Supplement Table Cell type markers). Using a 2-fold gene expression level difference between homozygous groups (i.e., |log_2_FC|>1), 964 QTT genes under additive eQTLs (53,157 eQTLs associated to 12,572 genes), 32 QTT genes under *X. maculatus* allele dominant eQTLs (61 eQTLs associated to 42 genes), and 5 QTT genes under *X. couchianus* allele dominant eQTLs (5 eQTLs associated to 4 genes) were used for further analyses (Supp Data-Qualified eQTLs-QTT). There are 14 QTT genes under additive eQTLs found to be over-enriched in different cell types. These genes include *wnk1b*, *popdc2* in cell type 5 oligodendrocyte (Fig. 4A), *sall3a* in cell type 9 microglial (Fig. 4B), *disc1* and PLEKHA4-like in cell type 10 astrocyte (Fig. 4C), *ano2b*, *hpca*, *scn1ba* and *nhsl1a* in cell type 11 neuron (Fig. 4D), and *cyp2n13*, *pdgfbb*, *rab11fip5a*, *fcho2*, and *slc2a1b* in cell type 13 endothelial (Fig. 4E).

**Figure 4.**
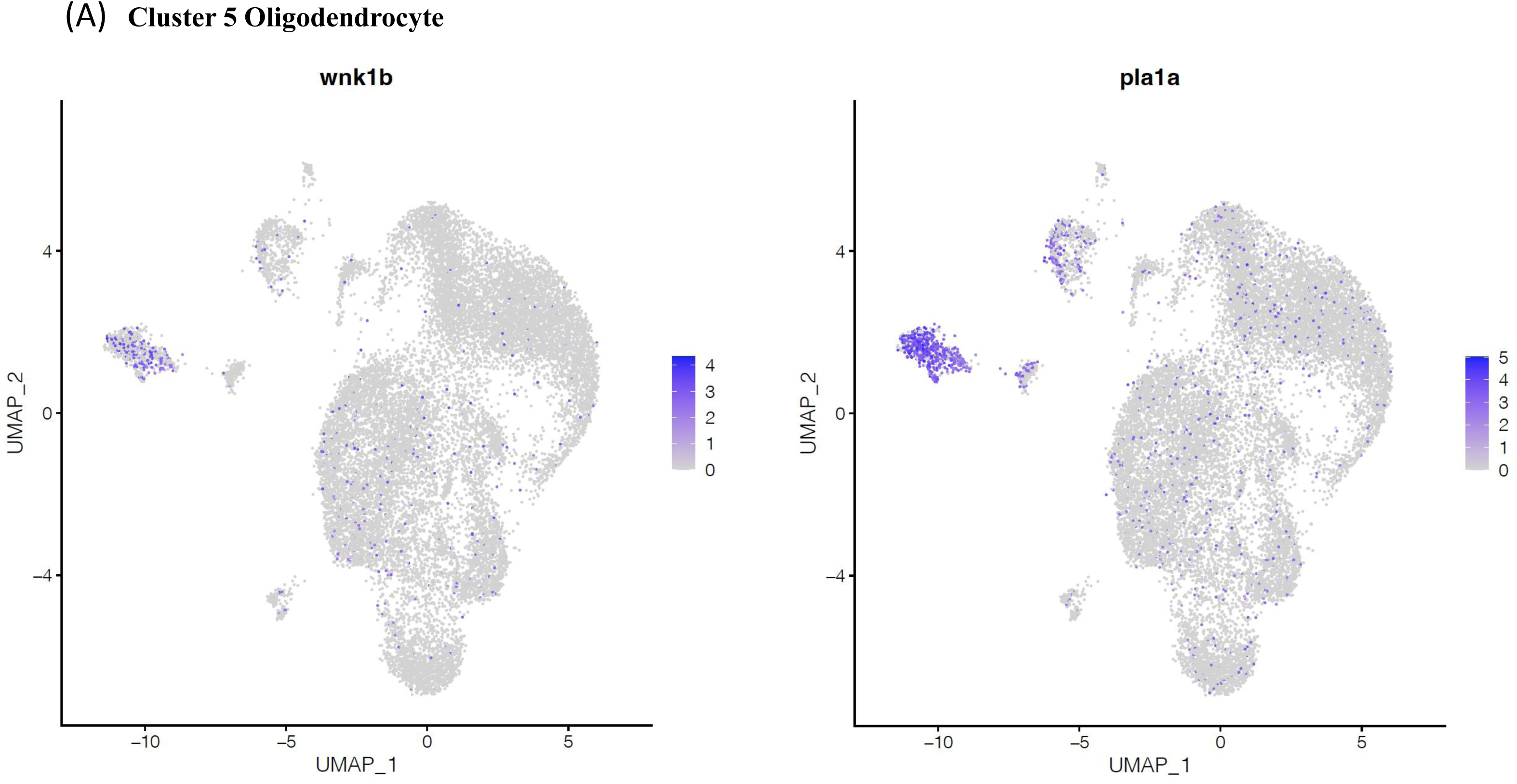
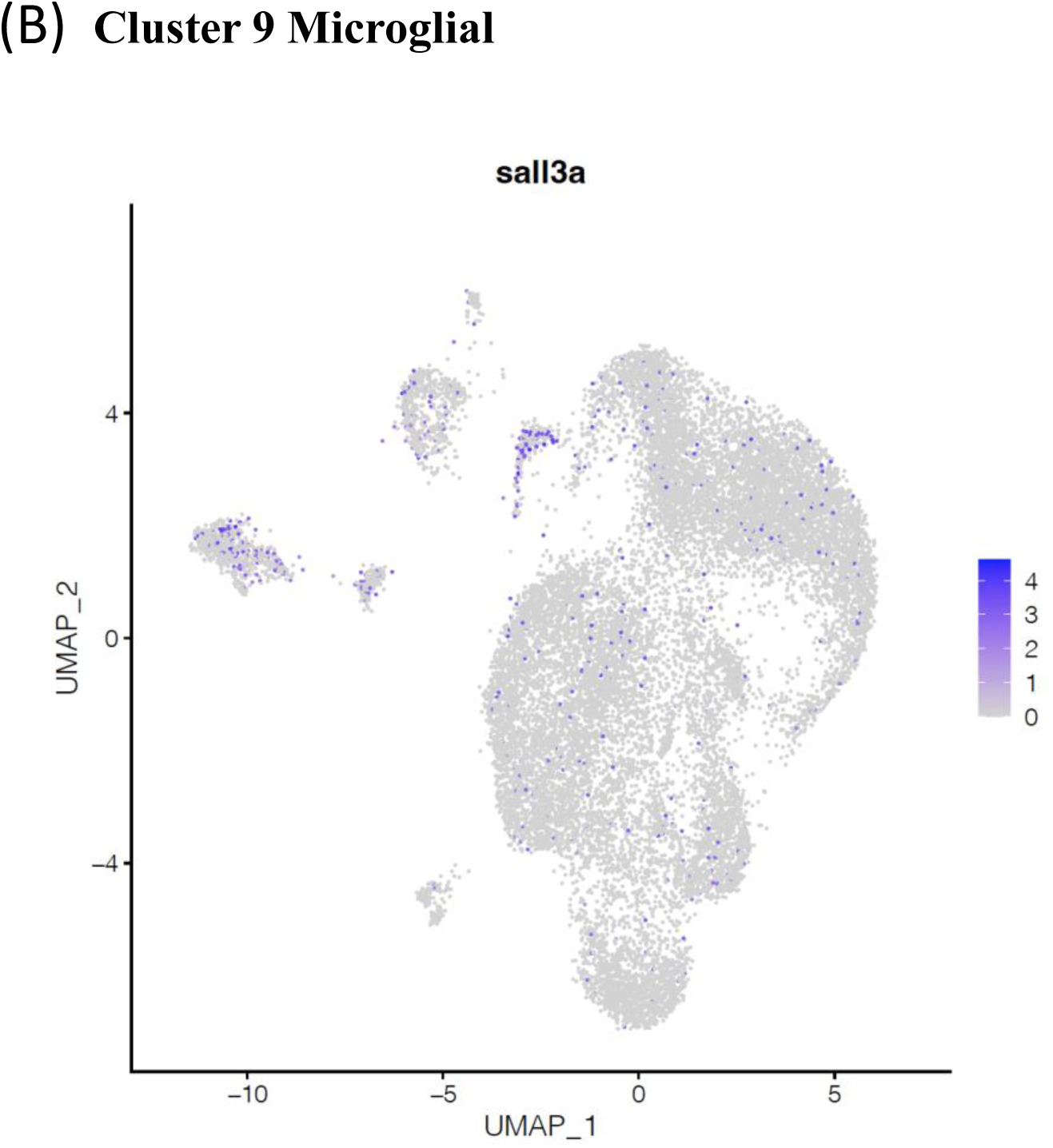
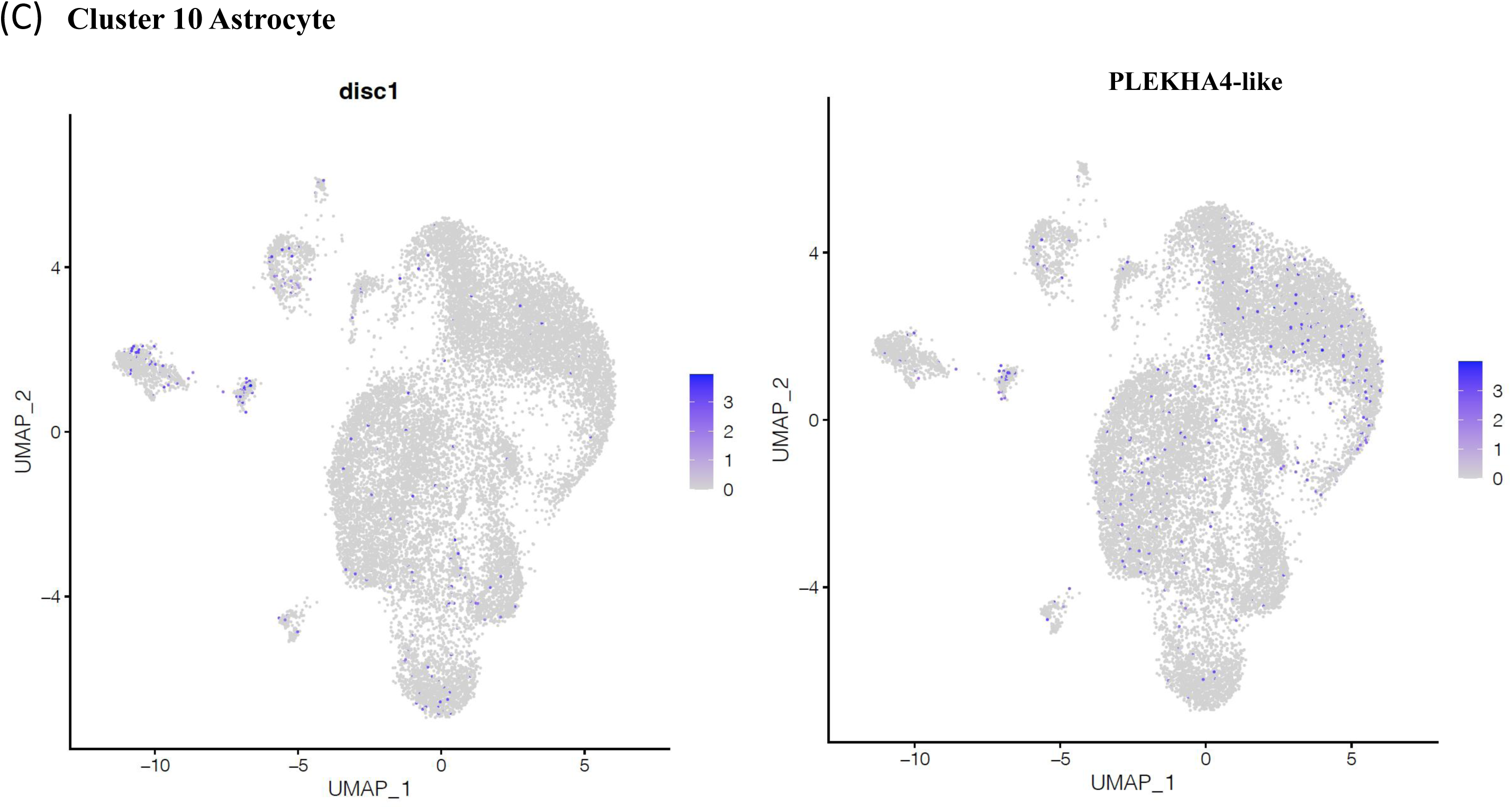
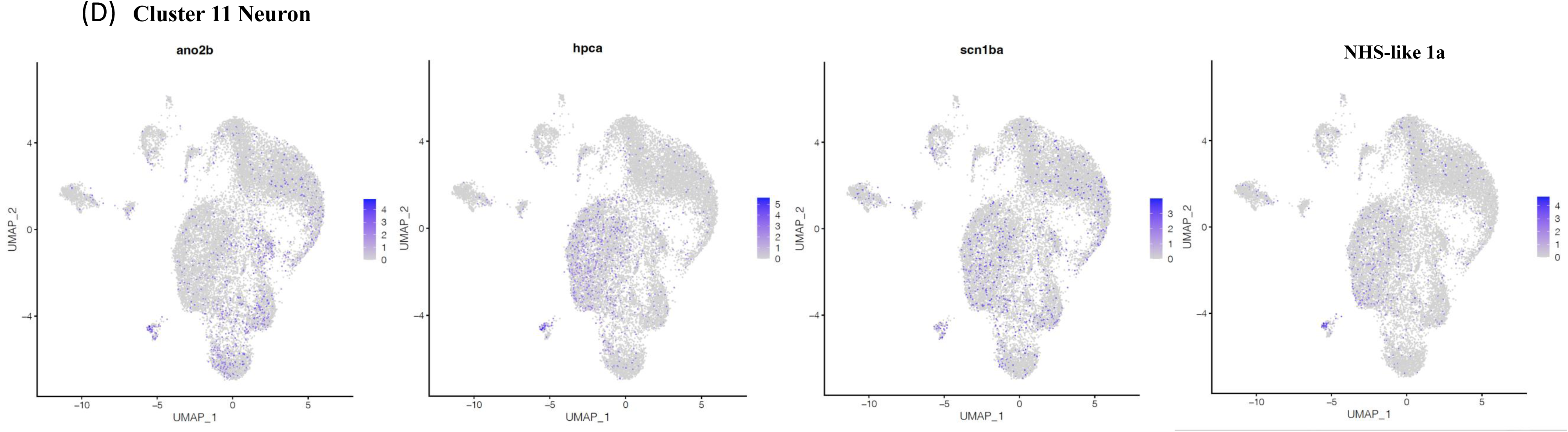
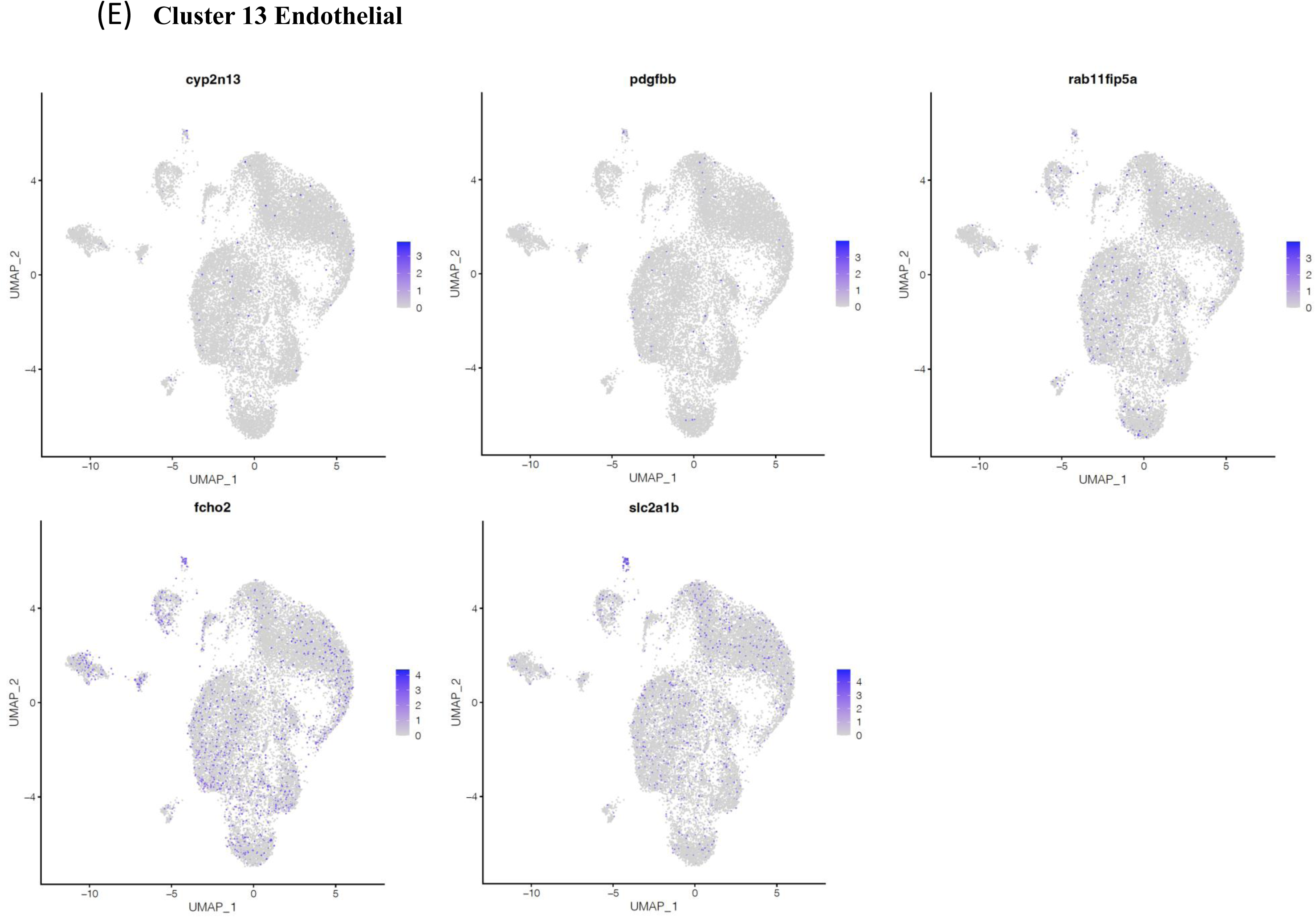
Cell type-specific QTT. Expression of a total of 14 QTT genes exhibited cell-type specific expression are plotted on UMAP. Subfigure A-E represent oligodendrocyte, microglial, astrocyte, neuron and endothelial cells. For each individual plot, dot on the UMAP represent a sequenced nucleus, and colors of dots represent normalized gene expression level of plotted gene.

## Discussion

In this study, we investigated interspecies hybrids between two distantly related *Xiphophorus* fish species with the main purpose to quantity and identity cell types exhibiting inter-specific genetics interactions. Extreme phenotypes (e.g., transgressive segregation) are usually the result of incompatible cis- and trans-regulations as observed in hybrid infertility (20). Earlier studies adopting the same parental species as this study produced F_1_ interspecies hybrids displaying molecular phenotypes in fin and skin that are out of the parental ranges (17).

However, there is an overall lack of over- and under-dominant eQTLs with effect size comparable to additive and dominant eQTLs despite using a lower statistical stringency (Fig. 1; Supp. Fig. 2 & 3). This observation indicates that extreme F_1_ phenotypes may depend on specific heterozygous combination of multiple alleles that are rarely maintained across all individuals in the segregating F_2_ population (21, 22). This will be tested in a future study with larger F_2_ population size.

Males of the two parental species exhibit mating behavioral differences (23)*. X. maculatus* males only mate with female belonging to the same species, while *X. couchianus* males indiscriminately mate with females regardless of their species. This behavior discrepancy may be adaptive as *X. maculatus* are sympatric to other species while *X. couchianus* are not (24). Our breeding records in producing interspecies hybrids show that breeding *X. couchianus* male with non-*couchianus* female could yield hybrids, while the other breeding direction does not result in broods unless through artificial insemination (Supp. Fig. 1). This supports the unique *X. couchianus* male mating behavior accounts for successful un-assisted interspecies breeding, and suggest their neural circuit involved in recognizing females belonging to the same species is diverged from *X. maculatus*. We found QTTs and eQTL related genes include known genes involved in *Xiphophorus* cognition capability (25). QTTs include cognition related genes *apba1*, *cntn1, disc1, magi2*, *ndufa2*, and *st8sia1* (25, 26). In addition, out of 45 known genes associated to human intelligence, 21 fish orthologs are identified in eQTL related genes (*wnt4, col16a1, nkiras1, rnf123, grk6, prr7, exoc4, apba1, jmjd1c, gbf1, eea1, dcaf5, sh2b1, zfhx3, dcc, cse1l, ndufa5, shank3, tcf20, bmpr2,* and *atxn2*; Supplement Table-Human intelligence genes.txt). The universal mating behavioral trait is *X. couchianus* dominant since male F_1_ interspecies hybrids also practice the universal courting behavior (Supp. Fig. 1). Two of the 5 QTTs under dominant *X. couchianus* eQTLs are *nostrin* and LOC111609877 (receptor-type tyrosine-protein phosphatase H-like, PTPRH-like; Supp. Fig. 5) showed no expression in *X. maculatus* homozygotes and are likely relevant to the universal courting behavior. The Nostrin gene is lowly expressed in brain. Its overexpression leads to redistribution and inhibition of nitric oxide (NO) release (27). Since NO is crucial in neurotransmission, blood flow regulation, and neuronal signaling, these functions may be reduced in *X. couchianus* and their F_1_ hyrbids due to higher *nostrin* expression. PTPRH is involved in brain immune response and potentially affect neuroinflammation. It has also been linked to neurodegenerative diseases. The higher expression of PTPRH-like gene in *X. couchianus* homozygotes and heterozygote may indicate that *X. couchianus* allele is associated to neuroinflammation and cognition differences. The behavioral differences between parentals and gene expression differences among different genotypes are not to conclude that *X. couchianus* are cognitively inferior. But the expression pattern of Nostrin and PTRPH-like may explain the organ level functional differences.

Finally, we traced 14 QTTs under additive eQTLs to specific cell types. This is not to conclude that only these 14 genes are cell type-specifically regulated as there are QTTs globally expressed in multiple cell types but rather to exemplify that genetic interactions can take place in a cell type-specific manner. Published studies confirm cell type-specific function of 10 genes, including *wnk1b*, *sall3a*, *disc1*, *ano2b*, *hpca*, *scn1ba*, *nhsl1a*, *pdgfbb*, *fcho2*, and *slc2a1b*. Despite there are only a handful of genes satisfying the statistical thresholds, it indicates there are functional differences between the two parental homozygotes. These differences can potentially be extrapolated to the two parental species in explaining observed behavioral divergence, and lead to new animal models for studying brain functions by practicing precision breeding over select eQTL or QTTs. The 10 QTT genes are involved in:

1. Regulation of ions, including Wnk1b as a regulator of Cl^-^ in oligodendrocyte, Ano2b as a calcium-activated chloride channel that controls neuronal spike frequency (28–30) and Scn1ba as a voltage-gated sodium channel (31). They are also involved in behaviors. For example, knock out of Ano2 in mice led to impairments in motor coordination and learning tasks (29), and Scn1b null mutation resulted in phenotypes including retarded growth, seizures, ataxia and death (31). In addition, several monoallelic variants in human SCN1B have been associated with febrile seizures, febrile seizures plus (FS+), early-onset absence epilepsy and focal epilepsy (32).
2. Neurogenesis, including Disc1 for dendrite development (33), Nhsl1a for lamellipodia stability and cell migration by stabilizing Scar/WAVE2, Arp2/3 complex activity (34), and Hpca for astrocyte differentiation (35). Disc1 is a well-known cognition related gene. Expression of Disc1 mutant in astrocytes has been shown to impairs adult neurogenesis and dendrite development of newborn neurons and cognition (33). Hpca-knockout in mice do not show any obvious structural abnormalities within the brain, but it led to deficits in spatial and associative memory (36, 37). In addition, aberrant activity of HPCA in human can increase vulnerability to oxidative stress-induced damage and result in neuronal death and neurodegeneration, and oxidative stress-induced brain diseases including stroke (38).
3. Blood vessel function and brain-blood barrier integrity. Relevant genes include Fcho mediating endocytosis, glucose transporter Slc2a1b (Glut1)(39), and endothelial cells secreted ligand Pdgfbb (40–43). Lowered level of GLUT1 arrests cerebral angiogenesis and causes Glut1 deficiency syndrome. Additionally, brain endothelial specific Glut1 depletion can trigger a severe neuroinflammatory response in the Glut1 deficient brain (39). The interaction between endothelial secreted and pericyte expressed Pdgfr-β leads to activation of signaling pathways regulation blood-brain barrier BBB integrity (40–43). Overall, lower expression of these above genes suggest *X. couchianus* may have a functionally different central neuron system.

In summary, we identified brain genetic interactions and characterized genes that are under regulations of cis- and trans-regulators, as well as brain cell types hosting these genetics interactions. The enrichment of neuronal disorders and disease related to genes under eQTL regulations suggest that *Xiphophorus* hybrid can develop neuronal disease due to inter-specific genetics interactions and can be investigated further to establish new animal models for research related to these conditions.

## Materials and Methods

### Animals

*X. maculatus, X. couchianus,* first-generation (F_1_) interspecies hybrids between *X. maculatus* and *X. couchianus*, and intercross hybrids (F_2_) used in this study were supplied by the *Xiphophorus* Genetic Stock (http://www.xiphophorus.txstate.edu/). The F_1_ interspecies hybrids between *X. maculatus* Jp163A strain fish and *X. couchianus* were produced by enforced mating using *X. maculatus* as the maternal parent, and F_2_ hybrids were produced naturally by male and female F_1_ hybrids. All fish were kept, and samples were taken in accordance with protocols approved by Texas State University IACUC (IACUC9048). All fish were dissected at the age of 9 months. At dissection, fish were anesthetized in an ice bath and upon loss of gill movement sacrificed by cranial resection. Brains were dissected into RNAlater (Ambion Inc.) and kept at -80 °C until use.

### RNA isolation

Brain samples were homogenized in TRI-reagent (Sigma Inc., St. Louis, MO, USA) followed by the addition of 200 μl/ml chloroform, vigorously shaken, and subjected to centrifugation at 12,000 g for 5min at 4°C. Total RNA was further purified using an RNeasy mini RNA isolation kit (Qiagen, Valencia, CA, USA). Column DNase digestion at 25°C for 15 min removed residual DNA. Each sample was independently maintained throughout the isolation process.

Total RNA concentration was determined using a Qubit 2.0 fluorometer (Life Technologies, Grand Island, NY, USA) and adjusted for sequencing library preparation. RNA quality was verified on an Agilent 2100 Bioanalyzer (Agilent Technologies, Santa Clara, CA) to confirm that RIN scores were above 7.0 prior to subsequent gene expression profiling.

### Inter-specific genetic variants identification and annotation

To identify interspecies polymorphisms between the *X. maculatus* and *X. couchianus*, whole genome shotgun sequencing reads of 4 *X. maculatus* and 5 *X. couchianus* were used (17). Raw sequencing reads were filtered using fastx_toolkit (http://hannonlab.cshl.edu/fastx_toolkit/index.html). Filtered sequencing reads were mapped to the reference *X. maculatus* genome (GenBank assembly accession: GCA_002775205.2) using Bowtie2 “head-to-head” mode (44).

Alignment files were sorted using Samtools (45). Pileup files were generated for each *X. maculatus*, and *X. couchianus* sample, and variant calling was processed by BCFtools for polymorphism detection, with minimum variant locus coverage of 2 and variant genotyping call Phred score of 0 and alternative genotyping Phred score ≥ 20 for BCFtools (45, 46). Inter-specific polymorphic sites are determined by variants where all sequenced *X. maculatus* samples support a homozygous *X. maculatus* reference base call, and *X. couchianus* samples support a homozygous alternative variant call. The inter-specific polymorphic sites were kept as a reference of genetic variance in BED format.

### Gene expression profiling

RNA sequencing was performed upon sequencing library construction using the Illumina TruSeq mRNA library preparation kit (Illumina, Inc., San Diego, CA, USA). RNA libraries were sequenced as 150bp paired-end fragments using Illumina NovaSeq system (Illumina, Inc., San Diego, CA, USA). Short sequencing reads were filtered using fastx_toolkit (47). RNA-Seq sequencing reads were produced from independent brain samples of F_2_ hybrids. Sequencing reads were mapped to *X. maculatus* reference genome (GenBank assembly accession: GCA_002775205.2) using Tophat2 (48). Gene expression was subsequently profiled by counting number of sequencing reads that mapped to gene models annotated by NCBI using Subread package FeatureCount function (49).

### Expression quantitative trait locus analyses

Sequencing adaptor contamination of brain-derived RNAseq reads was first removed from raw sequencing reads using fastx_toolkit, followed by trimming of low-quality sections of each sequencing read. Low quality sequencing reads were further removed from the sequencing result (http://hannonlab.cshl.edu/fastx_toolkit/index.html).

Processed sequencing reads were mapped to *X. maculatus* genome v5.0 (GenBank assembly accession: GCA_002775205.2) using Bowtie2 (44). Mpileup files were made using legend version of samtools (v1.13) and genotyping was processed using Bcftools (MAPQ >=30, Phred score of genotype call = 0, with alternative genotype call Phred score >=20). Genotype in this study refers to inheritance of ancestral alleles, with heterozygous meaning that a locus exhibited genetic material from both ancestors (i.e., *X. maculatus* and *X. couchianus*), and homozygous means that a locus exhibited genetic material from only the one parental species. Only genotypes of variant sites that are supported by reference inter-specific variants site were kept for further analyses. Gene specific expression counts were normalized to library size as counts per million reads (cpm). Per gene, a linear model was formed between the normalized gene expression values and copy numbers of alternative allele (i.e., *X. couchianus* allele) per variant site. Analysis of variance (ANOVA) was used to test gene expression differences among genotype groups. Genome wide multiple test correction was performed using Bonferroni method. Adjusted p-value < 0.05 and 2-fold expression differences between genotype groups per variant site were used to determine if a gene is under eQTL regulation.

### Single-nucleus RNA sequencing

Nuclei were isolated from *X. maculatus* brain using the Singulator2 instrument per manufacturers protocols (S2 Genomics; Livermore, CA). After mechanical disruption, cell filter straining steps, and washes, the cell nuclei were suspended in a nuclei isolation buffer (S2 Genomics; Livermore, CA), with all buffers containing 0.4U/ul RNAase inhibitor, prior to microfluidic encapsulation on the 10X Chromium instrument (10x Genomics®, Pleasanton, CA) to nanoliter-scale.

Single-nucleus libraries were generated using the GemCode Single-Cell Instrument and Single Cell 3′ Library and Gel Bead Kit v3 and Chip Kit (10x Genomics®, Pleasanton, CA) according to the manufacturer’s protocol. Before sequencing, every library was analyzed on a Bioanalyzer high-sensitivity chip to ensure the expected cDNA fragment size distribution was achieved. The appropriate number of individually barcoded GEM libraries were pooled and sequenced on a NovaSeq 6000 instrument (Illumina) with 2x150bp length using these sequencing parameters: 26 bp read 1 – 8 bp index 1 (i7) – 98 bp read 2 with 200 cycles.

The Cell Ranger software pipeline (version 3.1.0) was used to demultiplex cellular barcodes, map reads to the *X. maculatus* reference genome (GCA_002775205.2) and transcriptome using the STAR aligner, and to generate normalized aggregate data across samples, producing a matrix of gene counts versus cells. The individual tissue-specific sequenced Gel Bead-In Emulsion (GEM) libraries were each processed with the CellRanger v7.0.1 pipeline (10X Genomics) to create a cellular barcode by genomic feature matrix.

### Cell type annotation

Sequencing reads of snRNA-seq was mapped to *X. maculatus* reference genome (GCA_002775205.2) using ranger package (10X Genomics, Pleasanton, CA) with default setup of ‘count’ function. Mapping result, including gene expression count matrix, metadata, and reference gene names were forwarded to R package Seruet for cell cluster, cell cluster marker genes identification (50).

We annotated the *X. maculatus* brain cell clusters by comparing to annotated zebrafish, and mouse brain single cell (sc) RNAseq or snRNAseq datasets (51, 52). Cell cluster markers for each *X. maculatus* brain cell clusters were compared to marker genes of each annotated cell types of the reference datasets and tested for over-enrichment using hypergeometric test (p<0.05).

QTT associated genes were compared to cell type markers (avg_log2FC>0, p_val_adj <0.01, pct.1/pct.2>5, and pct.1>0.2). QTTs that are also cell type markers are determined to be cell type-specific QTT.

## Acknowledgments

This work was supported by the National Institutes of Health R24 award OD-031467 from the NIH Division of Comparative Medicine.

## Supplemental Figures

**Supplemental Figure 1.**
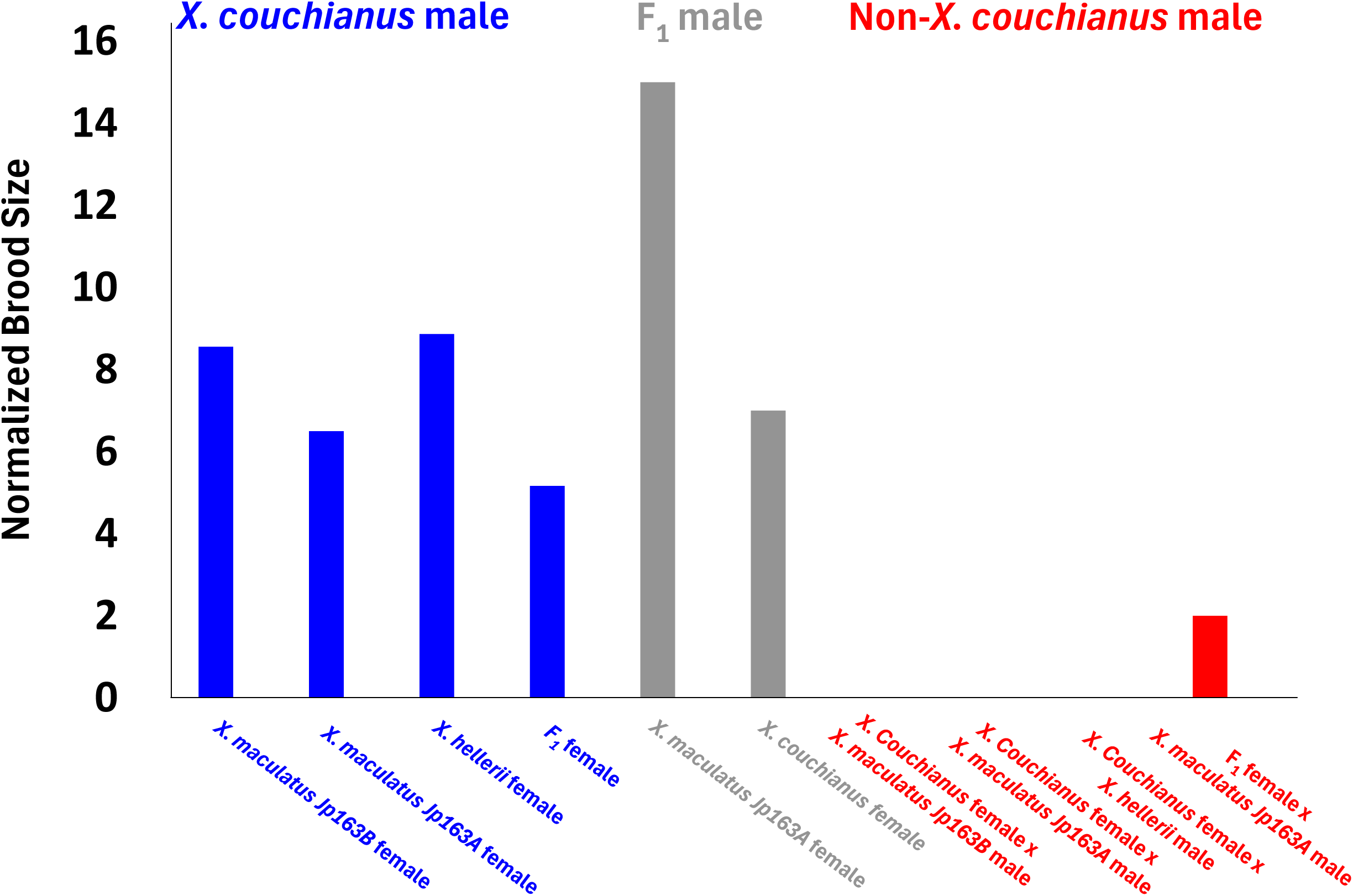
Breeding records of *X. couchianus* related crosses. Normalized brood sizes were calculated as total brood sizes divide by brood numbers for each cross. When male *X. couchianus* or F_1_ hybrids between *X. maculatus* and *X. couchianus* were used for breeding, broods were made regardless of the female species. However, when non-*X. couchianus* males were used to mate with *X. couchianus* female, no brood was made.

**Supplemental Figure 2.**
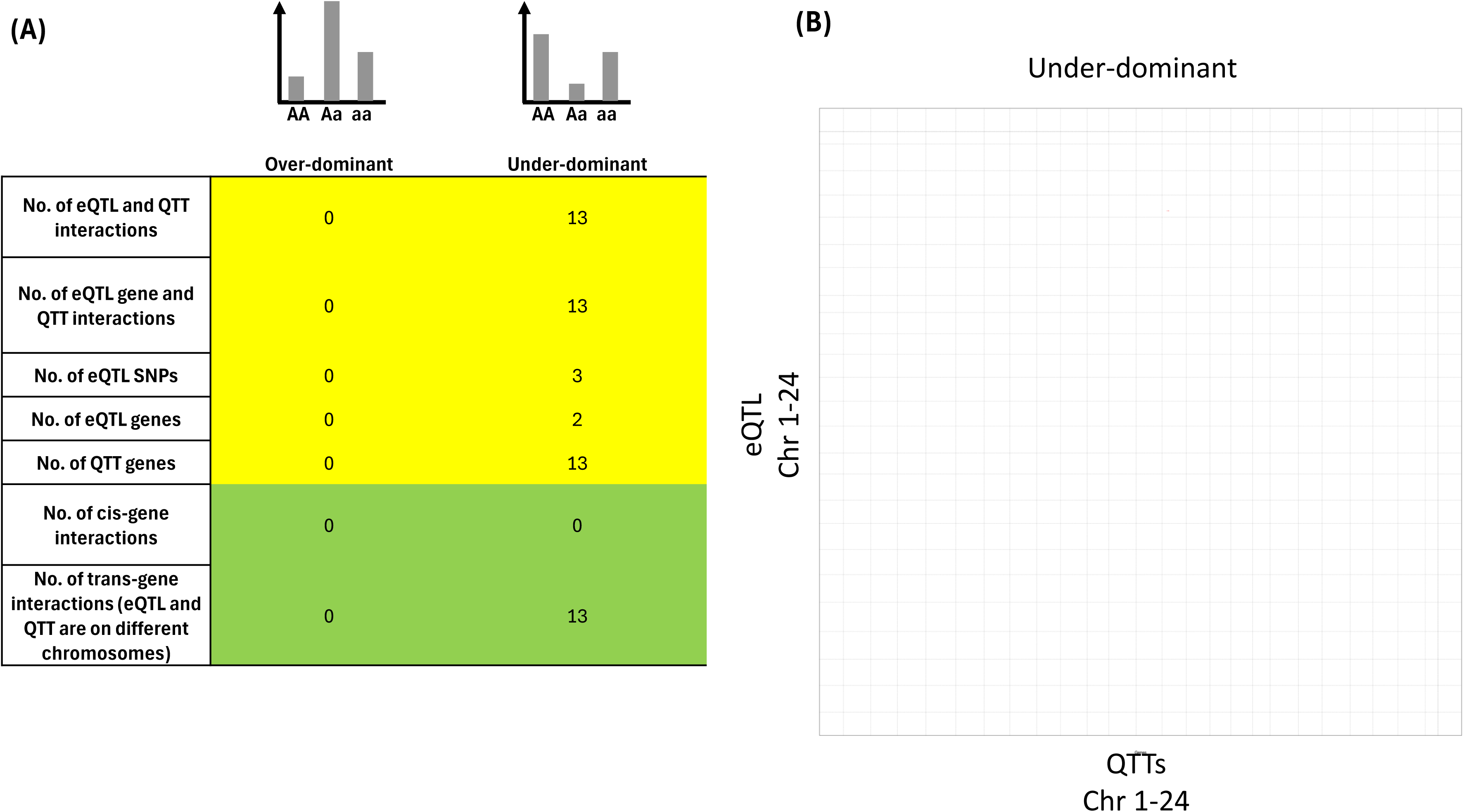
Quantification of over- and under-dominant eQTL and QTT. (A) Under-dominant eQTL were identified using FDR adjusted ANNOVA -log_10_p-value > 2. Yellow colored table reports quantity of eQTL-QTT interactions at inter-species polymorphism and gene levels. Green colored table reports quantity of cis- and trans-genetic interactions of each type. (B) Scatter plot shows genomic coordinates of under-dominant eQTLs and QTT genes. It is showed that heterozygosity of polymorphisms on chromosome 21 is associated with lower gene expression on chromosome 13 in heterozygote hybrids.

**Supplemental Figure 3.**
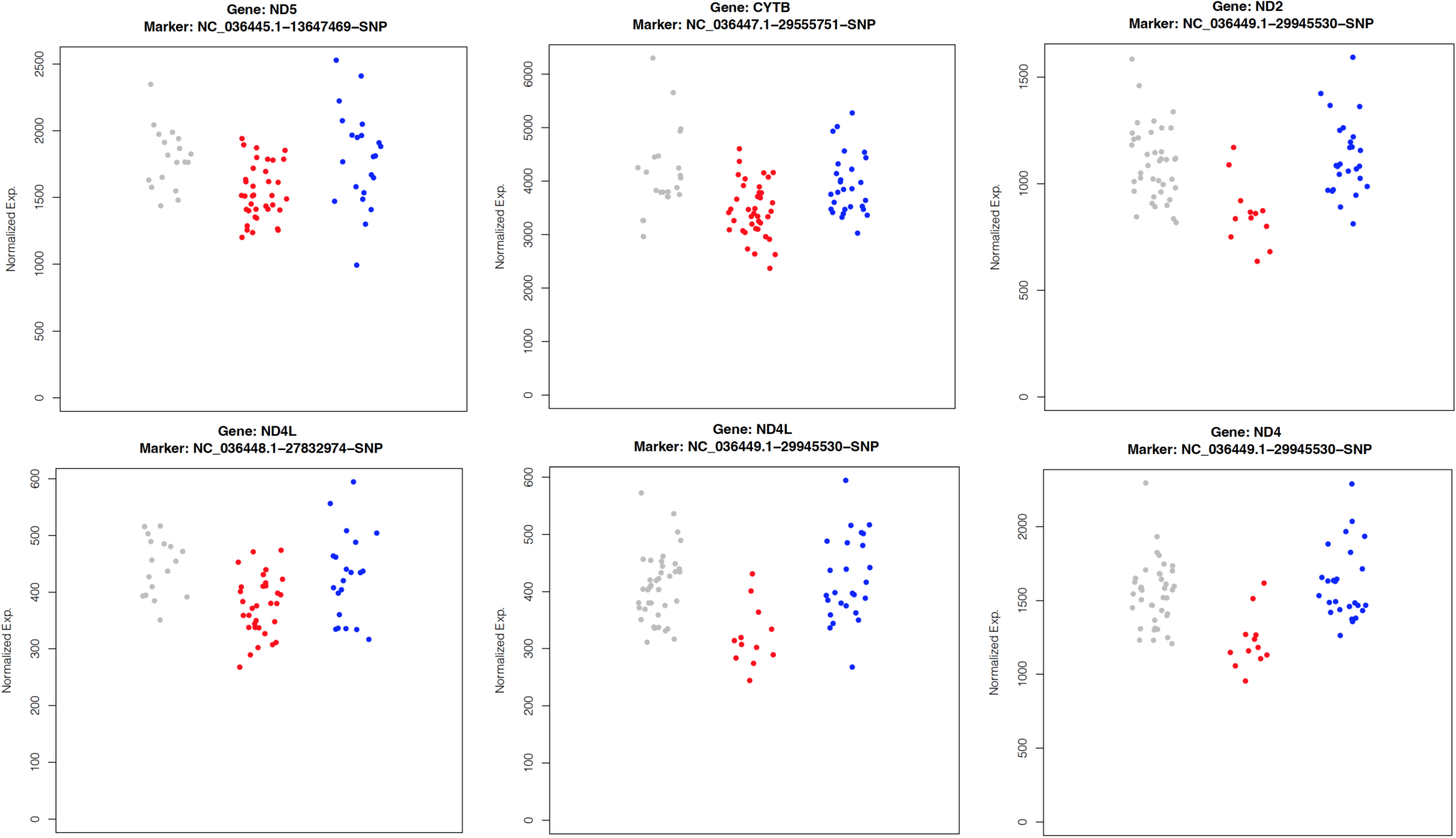
Under-dominant mitochondrial gene. Under-dominant QTTs encoded on mitochondrial DNA were identified using FDR adjusted ANNOVA -log_10_p-value > 2. For each plot, dot Y-axis coordinate represents library size normalized gene expression (See Materials and Methods), and color represents eQTL polymorphism genotype: gray, red and blue mean homozygous for P1, *X. maculatus*, allele; heterozygous for both parental alleles, and homozygous for P2, *X. couchianus* allele.

**Supplemental Figure 4.**
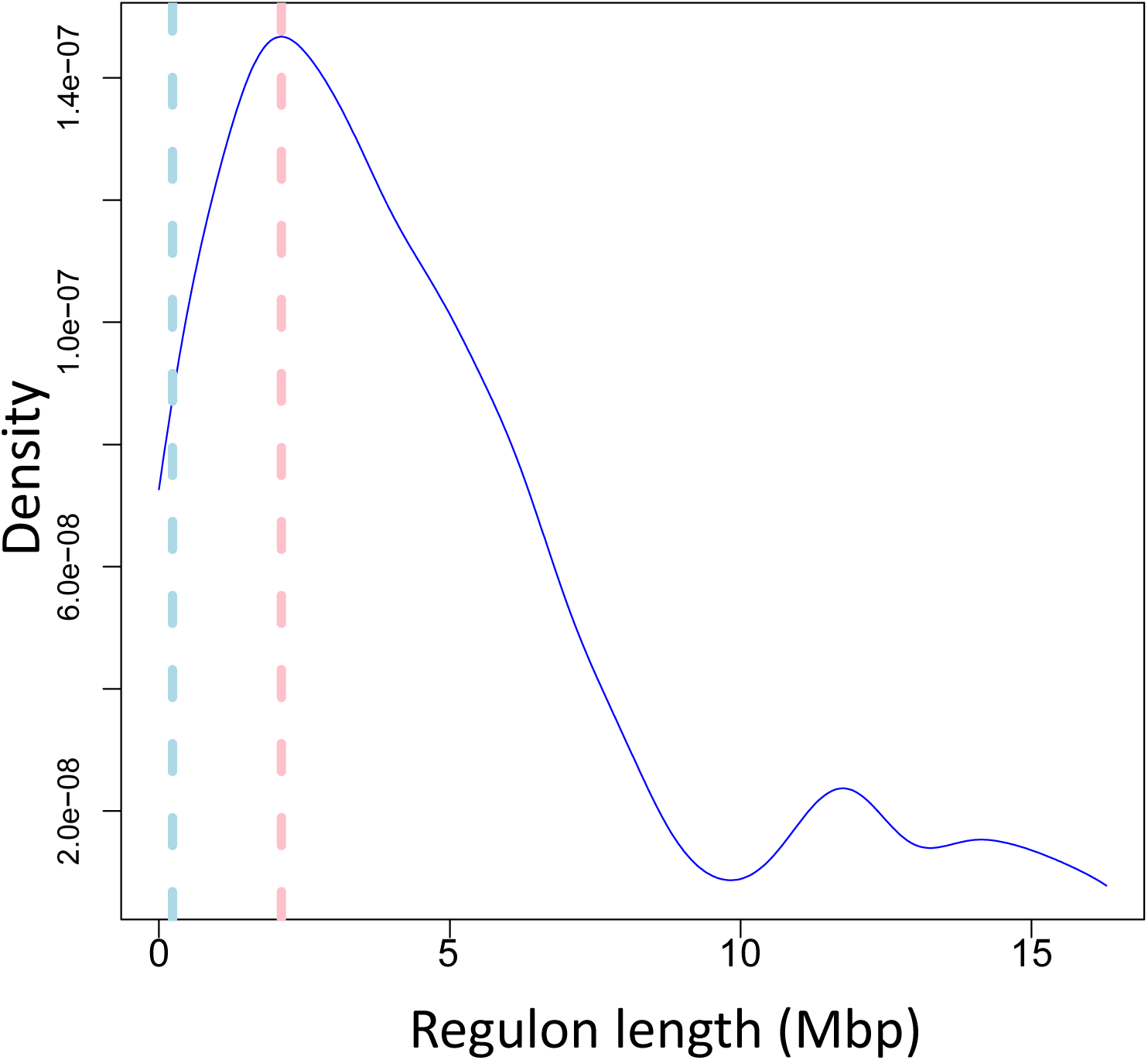
Density plot of additive eQTL distance from associated QTT. Distances between the most common cis- additive eQTLs and QTT gene were calculated and plotted as density plot. The X coordinate means distance between eQTL and QTT, and Y coordinate means percentage. Therefore, the area under curve is 1.

**Supplemental Figure 5.**
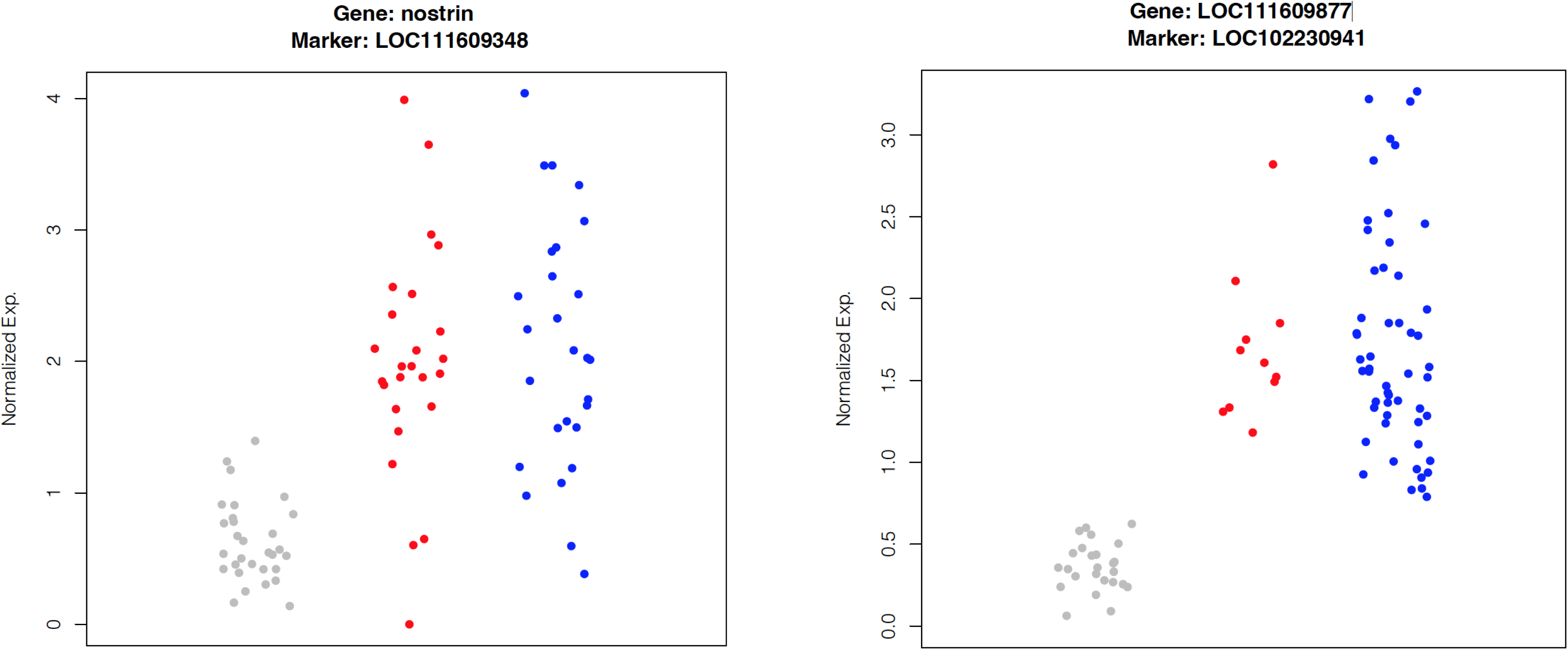
QTTs shown *X. couchianus* allele-dominant expression pattern. QTTs under *X. couchianus* allele dominant eQTL regulations were identified using FDR adjusted ANNOVA -log_10_p-value > 10, with log_2_FC>1 between heterozygotes and *X. maculatus* allele homozygotes. For each plot, dot Y-axis coordinate represents library size normalized gene expression (See Materials and Methods), and color represents eQTL polymorphism genotype: gray, red and blue mean homozygous for P1, *X. maculatus*, allele; heterozygous for both parental alleles, and homozygous for P2, *X. couchianus* allele.

## Supplemental Tables

**Supplemental Table 1. QTT Over-enriched functions**

**Supplemental Table 2. Human intelligence related genes**

**Supplemental Table 3. Cell type annotation**

**Supplemental Table 4. Cell type marker genes**

**Supplemental Data-MT.genotype**

**Supplemental Data-Qualified eQTL and QTT**

## Notes

### Competing Interest Statement

The authors have declared no competing interest.

