## Supplementary material for "Hybridization Reveals Cell Type-Specific Regulatory Variation Driving Brain Transcriptomic Divergence": Supp. Figs

### Slide 1
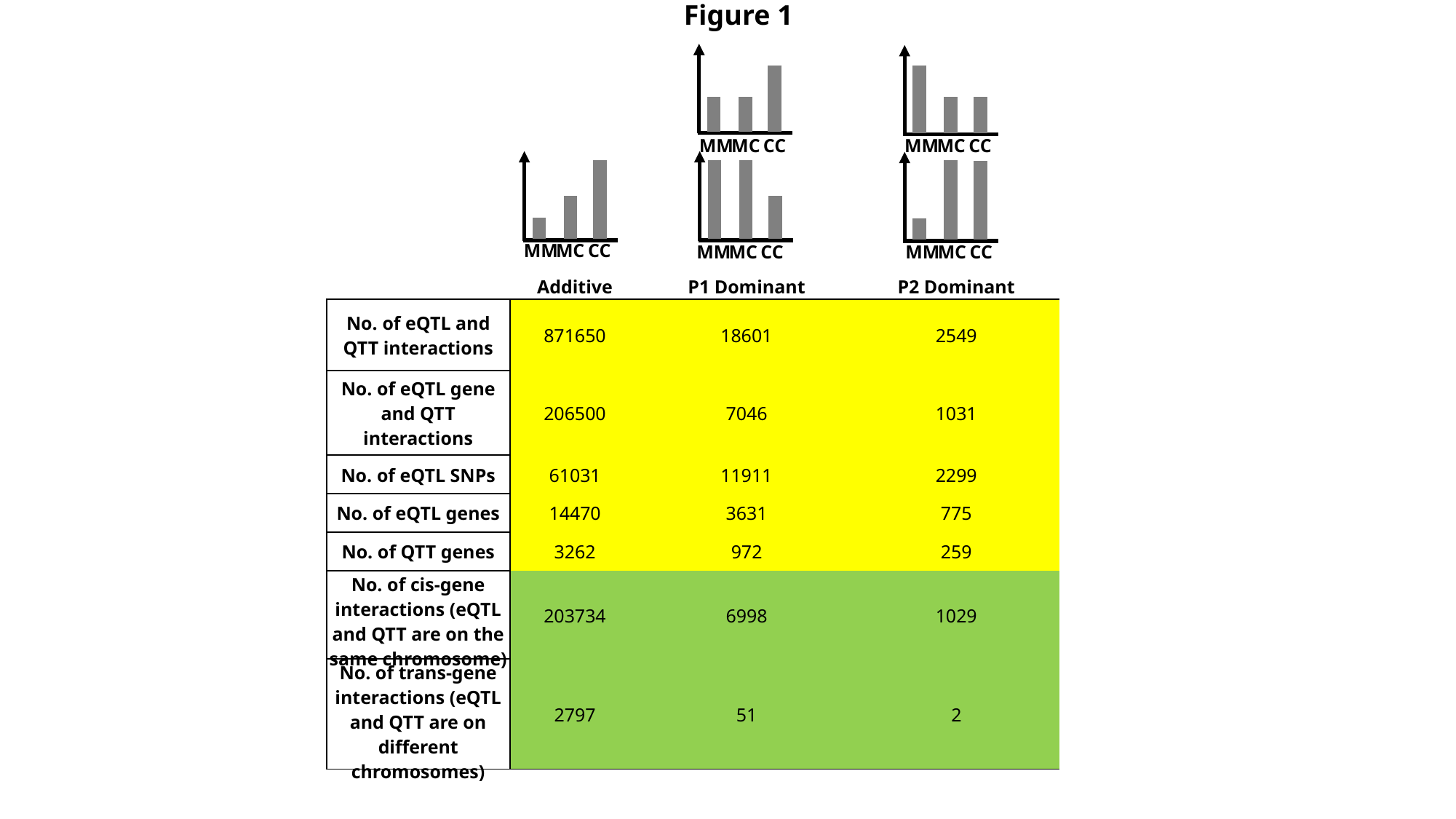

Figure 1
MM
MC
CC
MM
MC
CC
MM
MC
CC
MM
MC
CC
MM
MC
CC
| | Additive | P1 Dominant | P2 Dominant |
| --- | --- | --- | --- |
| No. of eQTL and QTT interactions | 871650 | 18601 | 2549 |
| No. of eQTL gene and QTT interactions | 206500 | 7046 | 1031 |
| No. of eQTL SNPs | 61031 | 11911 | 2299 |
| No. of eQTL genes | 14470 | 3631 | 775 |
| No. of QTT genes | 3262 | 972 | 259 |
| No. of cis-gene interactions (eQTL and QTT are on the same chromosome) | 203734 | 6998 | 1029 |
| No. of trans-gene interactions (eQTL and QTT are on different chromosomes) | 2797 | 51 | 2 |

### Slide 2
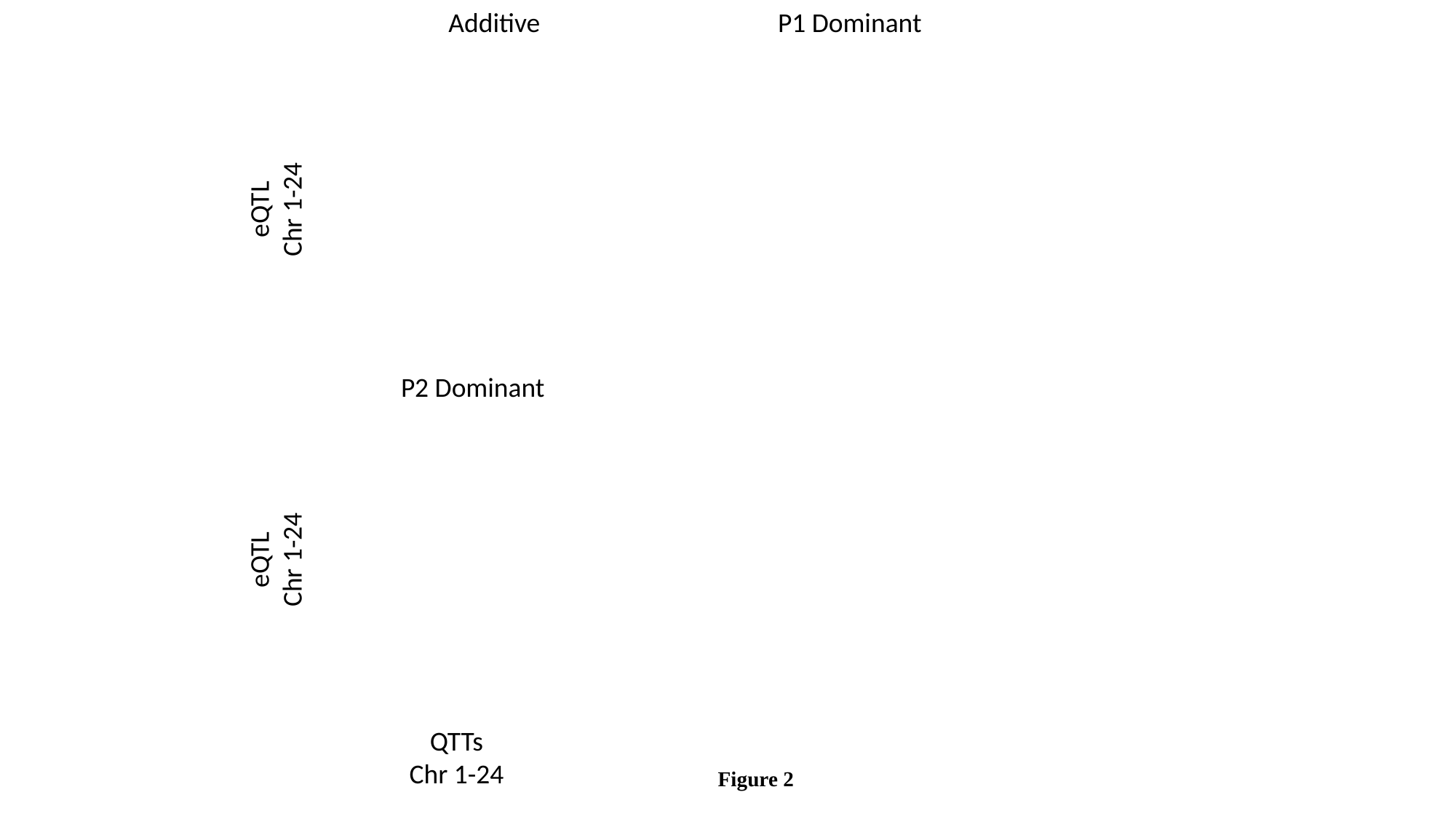

Additive
P1 Dominant
eQTL
Chr 1-24
P2 Dominant
eQTL
Chr 1-24
QTTs
Chr 1-24
Figure 2

### Slide 3
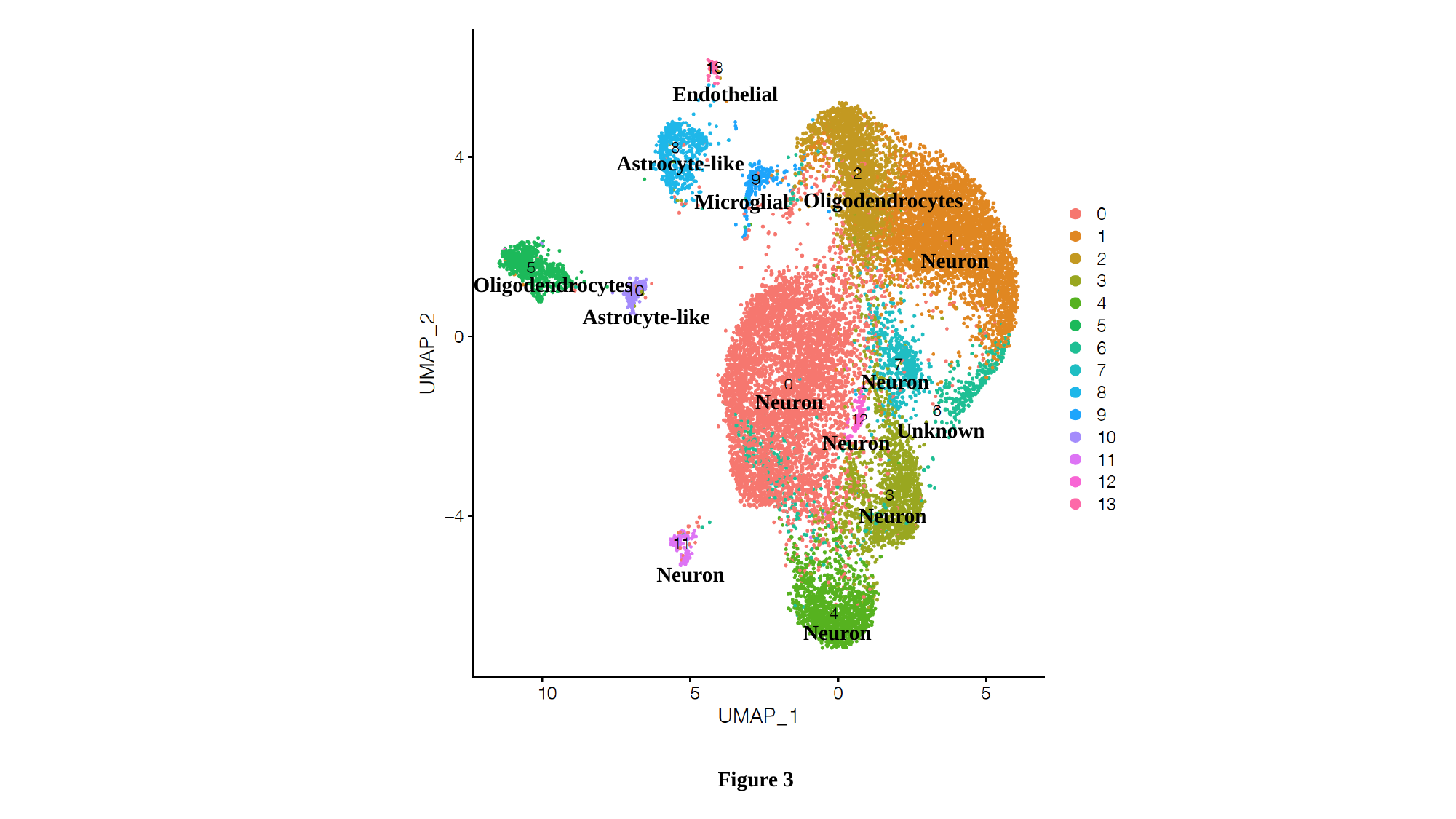

Endothelial
Astrocyte-like
Oligodendrocytes
Microglial
Neuron
Oligodendrocytes
Astrocyte-like
Neuron
Neuron
Unknown
Neuron
Neuron
Neuron
Neuron
Figure 3

### Slide 4
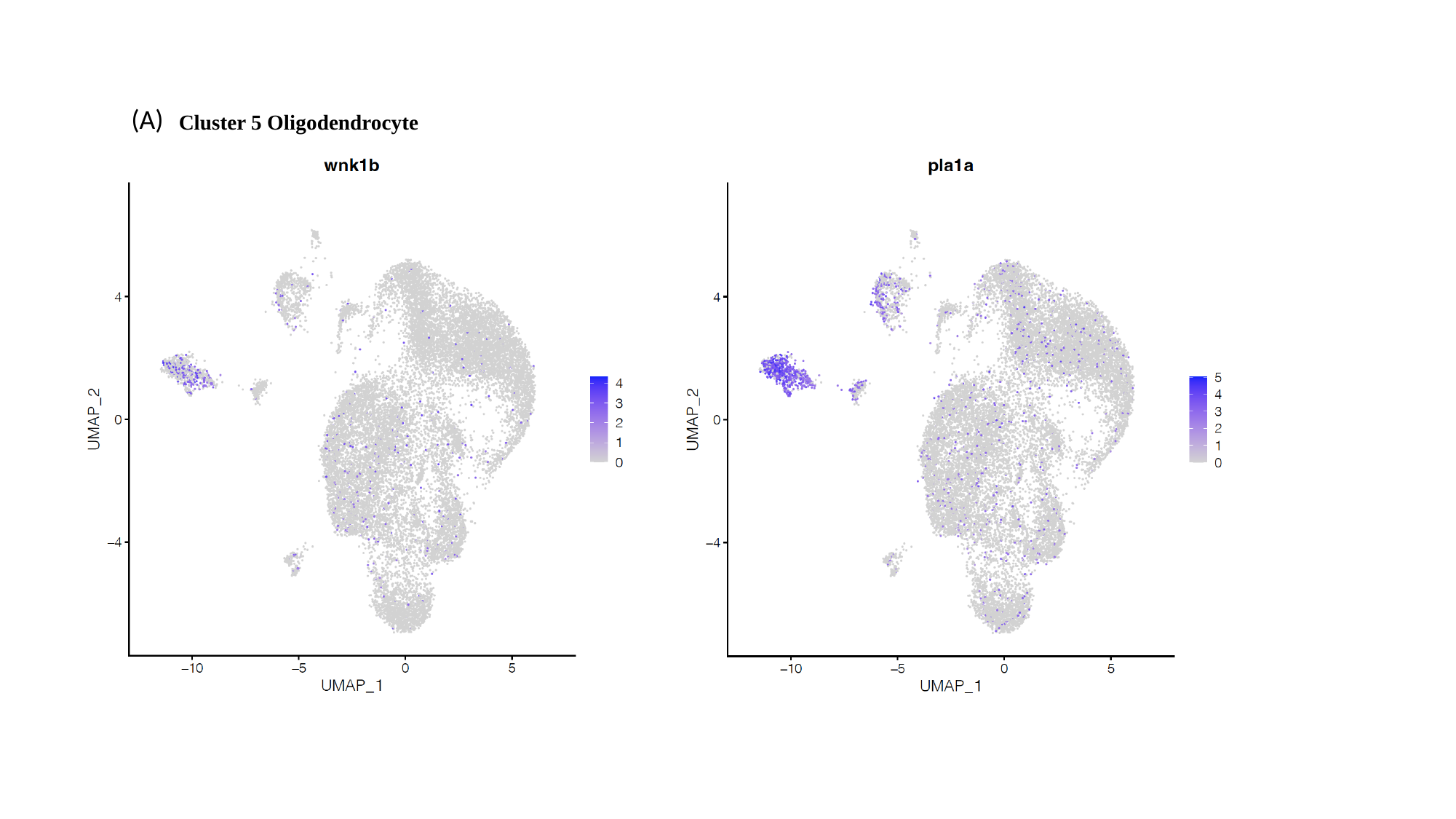

(A)
Cluster 5 Oligodendrocyte

### Slide 5
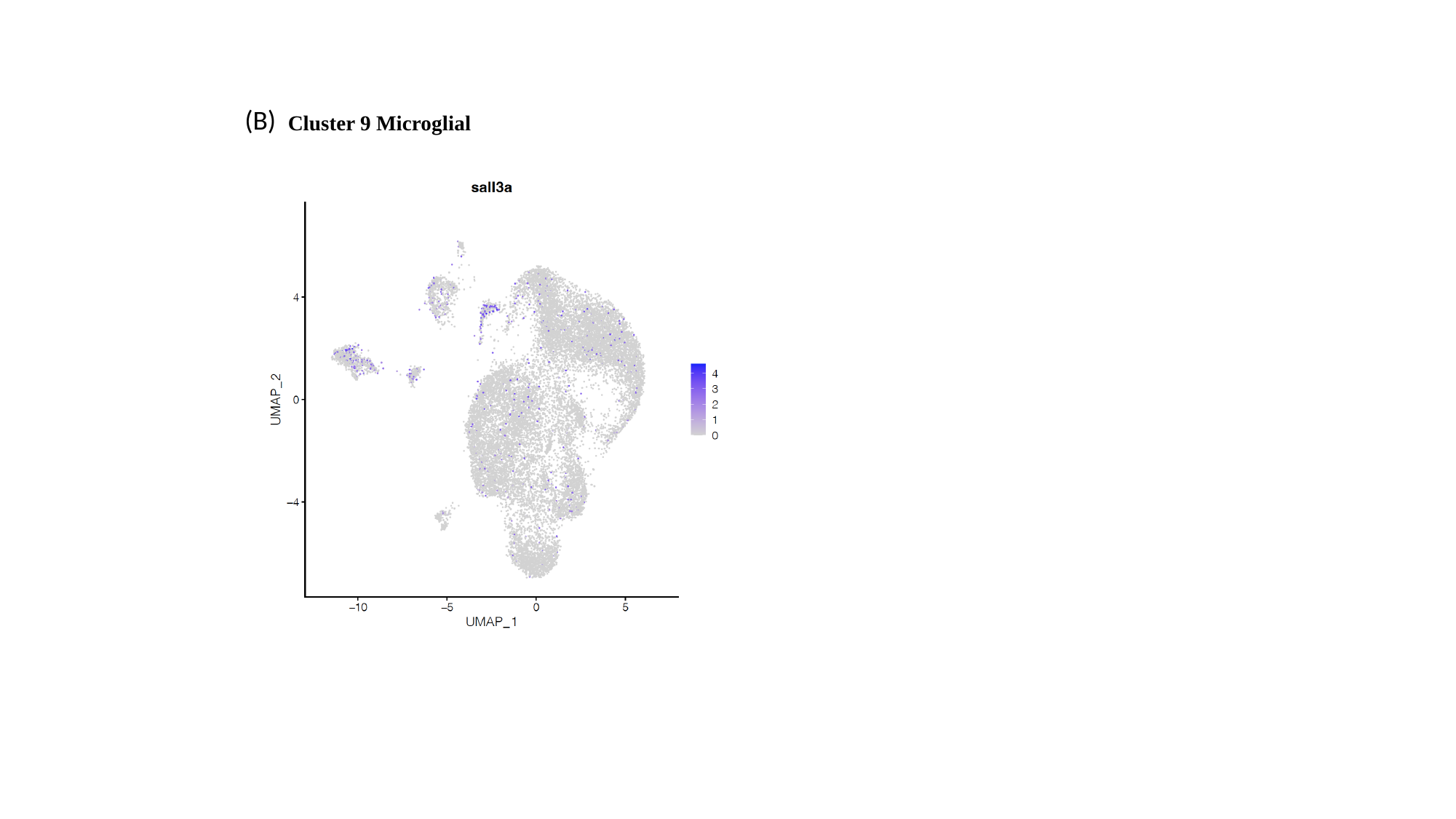

(B)
Cluster 9 Microglial

### Slide 6
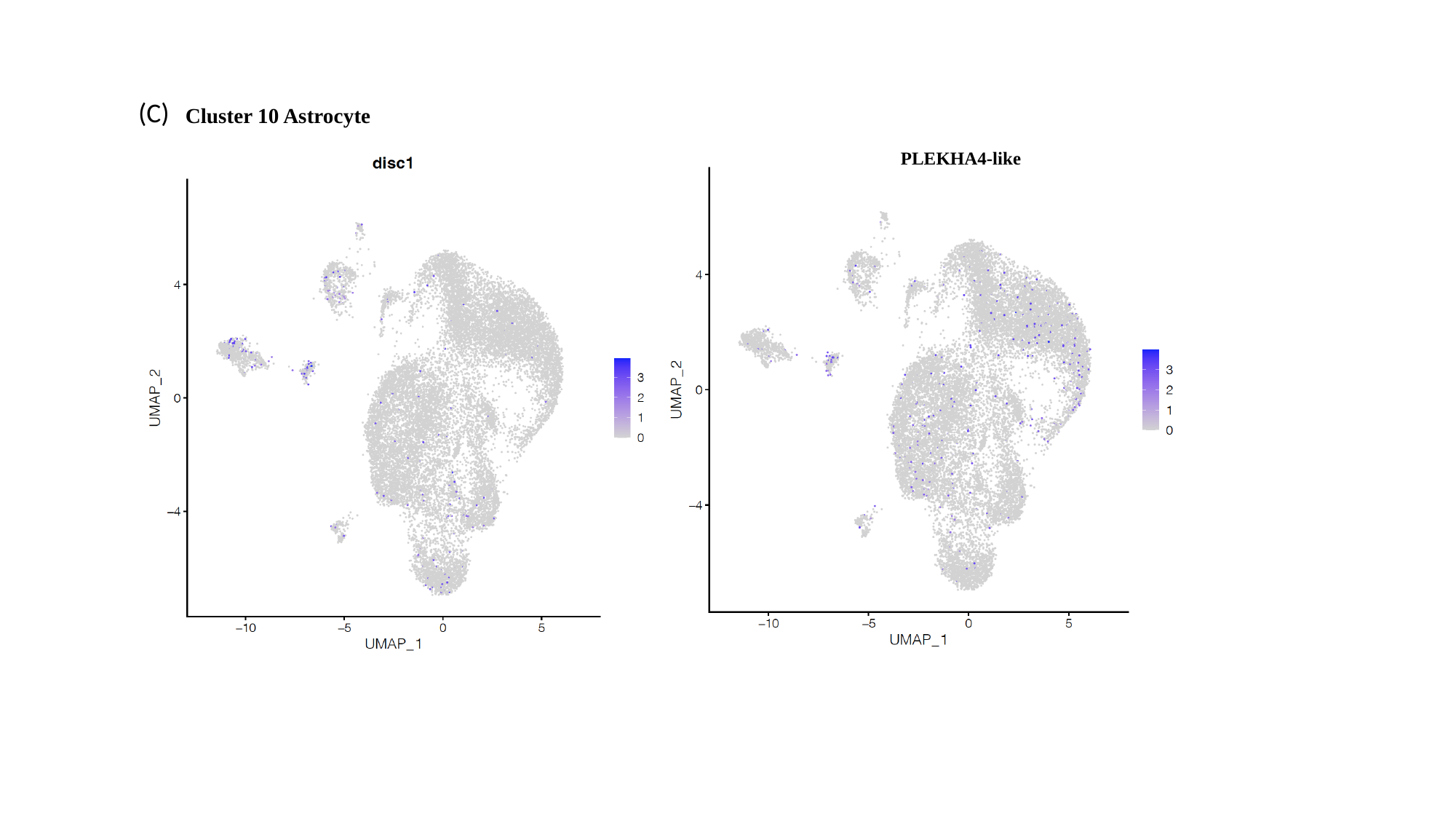

(C)
Cluster 10 Astrocyte
PLEKHA4-like

### Slide 7
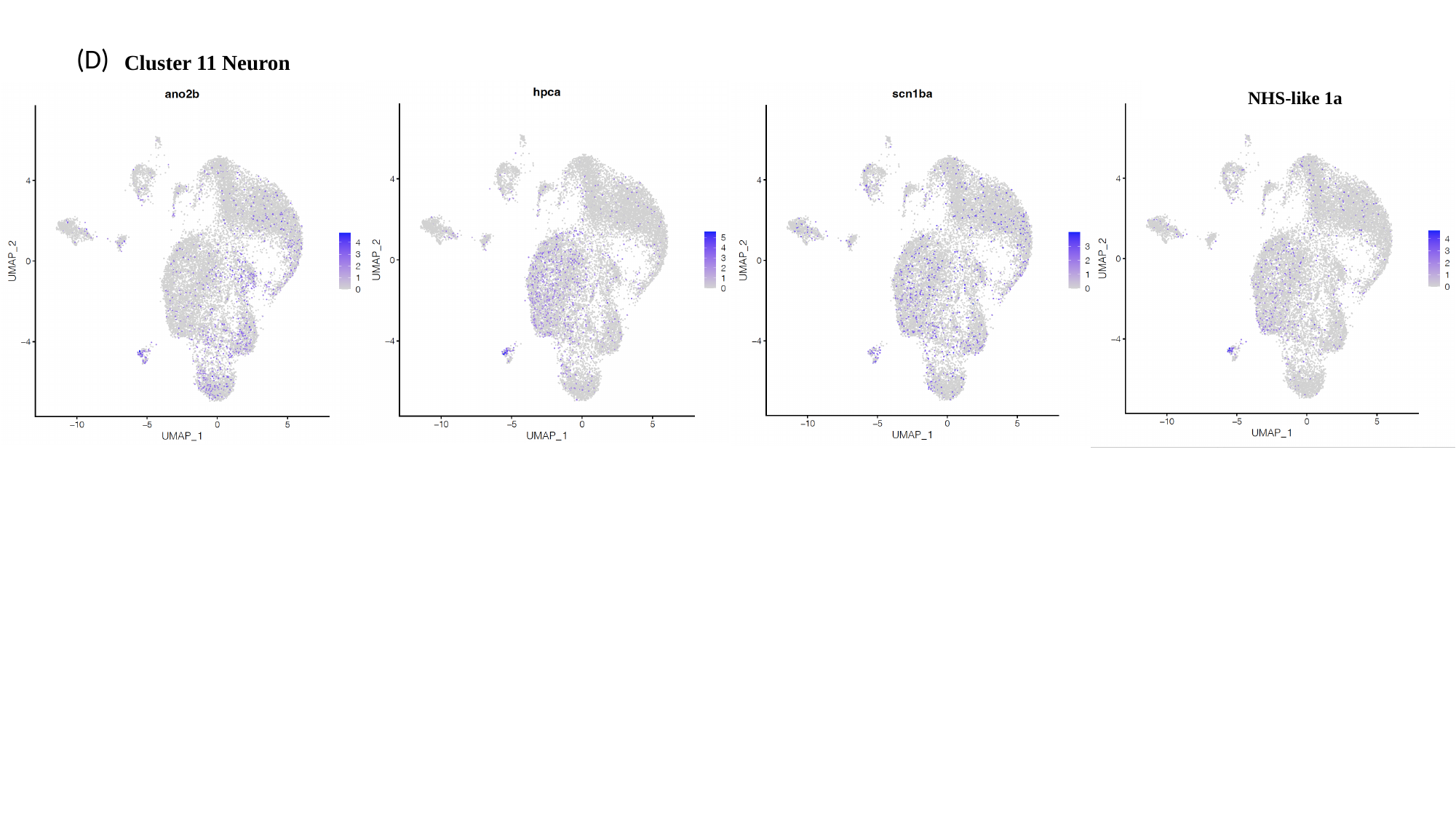

(D)
Cluster 11 Neuron
NHS-like 1a

### Slide 8
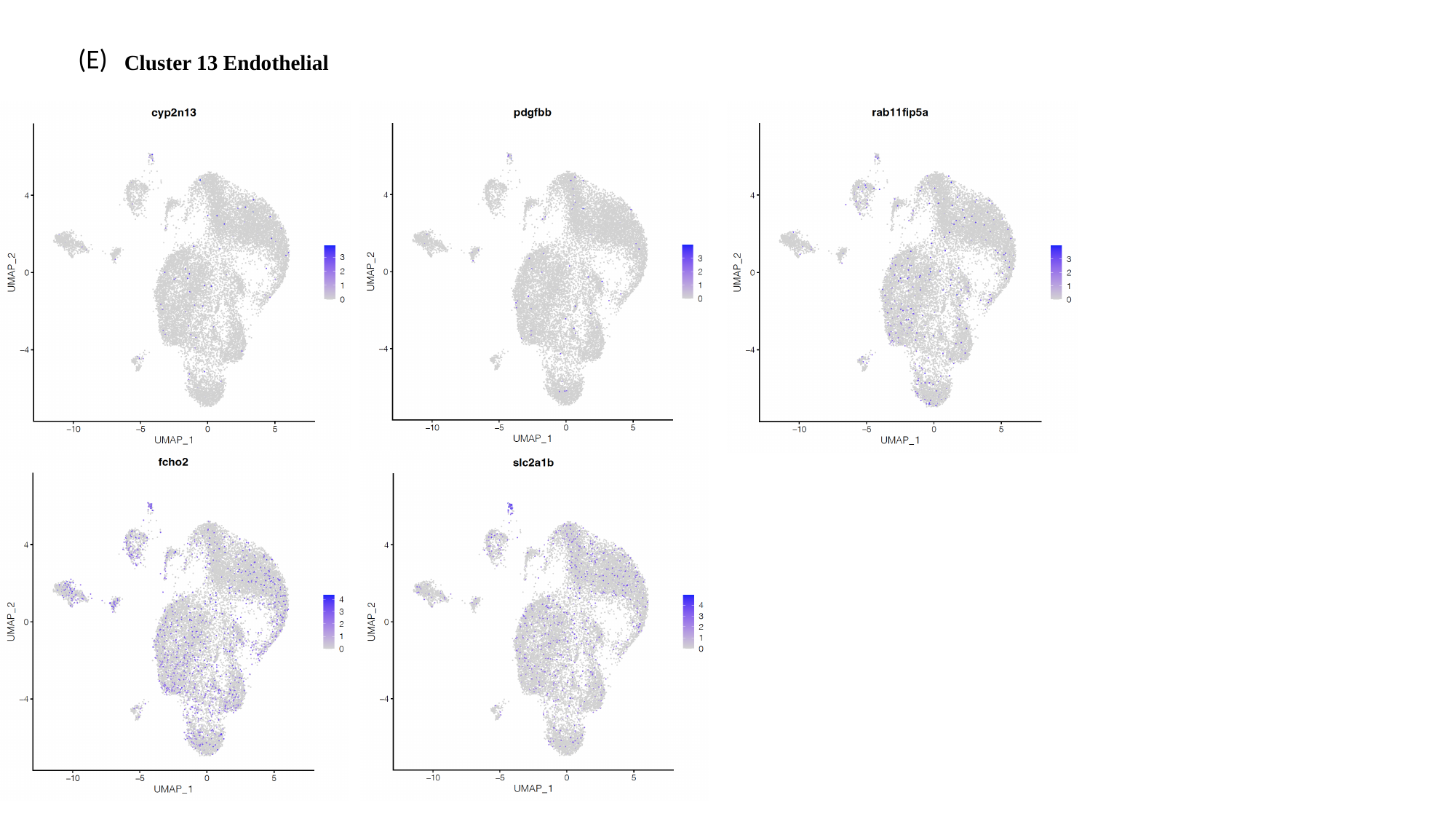

(E)
Cluster 13 Endothelial

### Slide 9
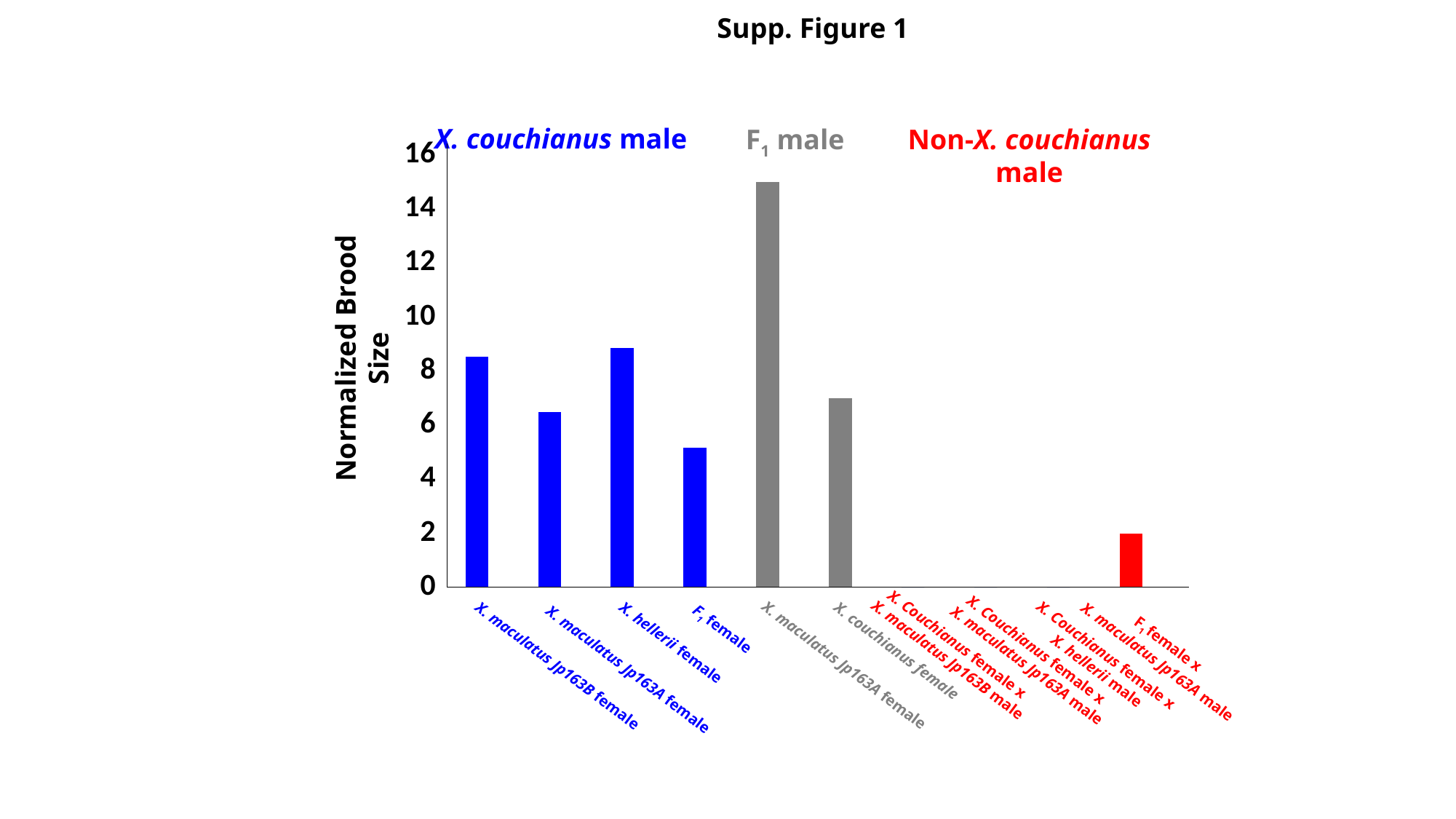

Supp. Figure 1
#### Chart
| Category | |
|---|---|
| X. maculatus jp163B F x X. couchianus M | 8.555555555555555 |
| X. maculatus jp163A F x X. couchianus M | 6.5 |
| X. hellerii F x X. couchianus M | 8.866666666666667 |
| F1 F x X. couchianus M | 5.166666666666667 |
| X. maculatus jp163A F x F1 M | 15.0 |
| X. couchianus F x F1 M | 7.0 |
| X. couchianus F x X. maculatus jp163B M | 0.0 |
| X. couchianus F x X. maculatus jp163A M | 0.0 |
| X. couchianus F x X. hellerii M | 0.0 |
| F1 F x X. maculatus jp163A M | 2.0 |X. couchianus male
F_1_ male
Non-X. couchianus male
Normalized Brood Size
X. Couchianus female x
X. maculatus Jp163B male
F_1_ female x
X. maculatus Jp163A male
X. Couchianus female x
X. maculatus Jp163A male
X. Couchianus female x
X. hellerii male
X. maculatus Jp163A female
X. couchianus female
X. hellerii female
X. maculatus Jp163B female
F_1_ female
X. maculatus Jp163A female

### Slide 10
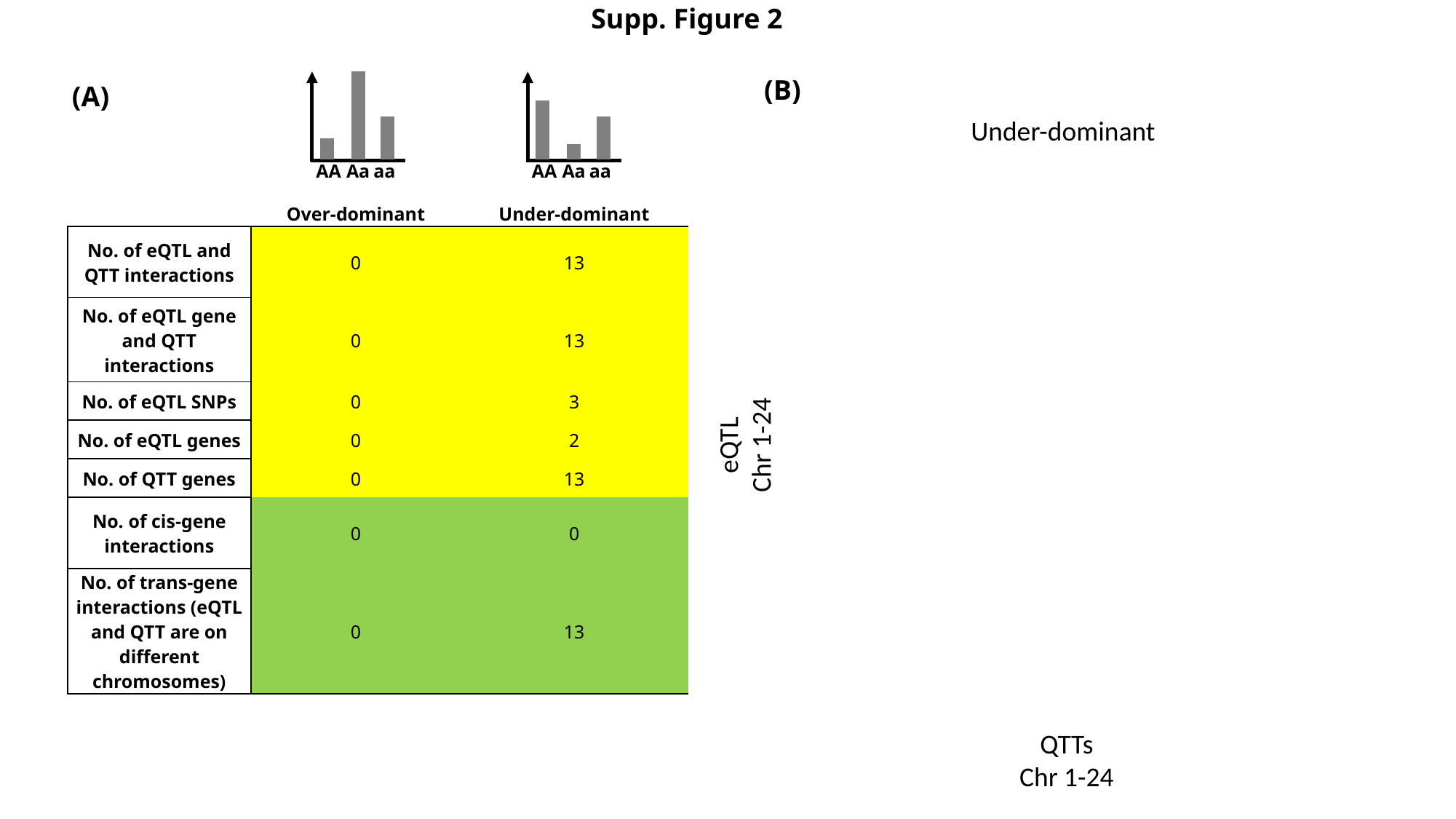

Supp. Figure 2
(B)
(A)
Under-dominant
AA
Aa
aa
AA
Aa
aa
| | Over-dominant | Under-dominant |
| --- | --- | --- |
| No. of eQTL and QTT interactions | 0 | 13 |
| No. of eQTL gene and QTT interactions | 0 | 13 |
| No. of eQTL SNPs | 0 | 3 |
| No. of eQTL genes | 0 | 2 |
| No. of QTT genes | 0 | 13 |
| No. of cis-gene interactions | 0 | 0 |
| No. of trans-gene interactions (eQTL and QTT are on different chromosomes) | 0 | 13 |
eQTL
Chr 1-24
QTTs
Chr 1-24

### Slide 11
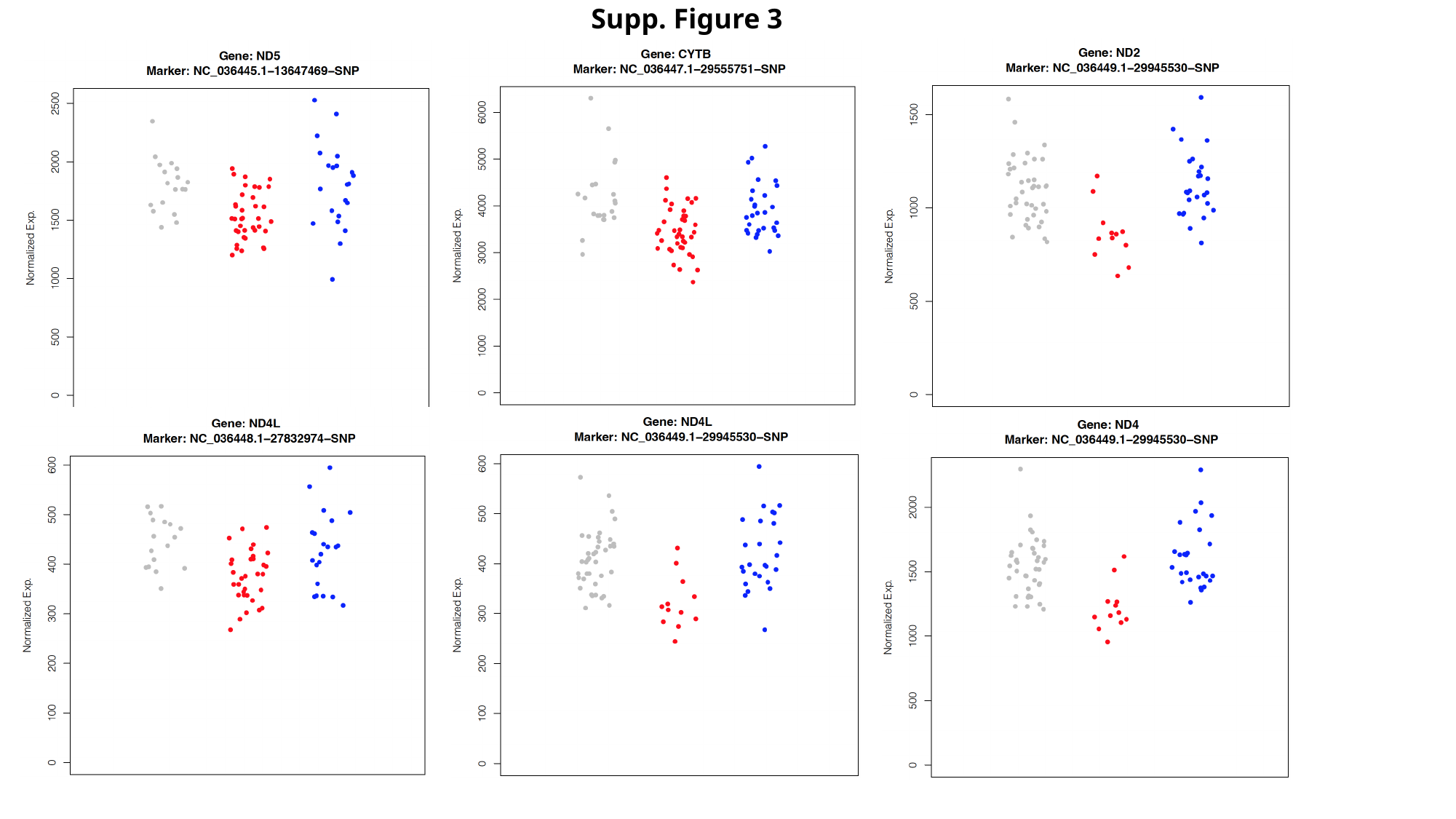

Supp. Figure 3

### Slide 12
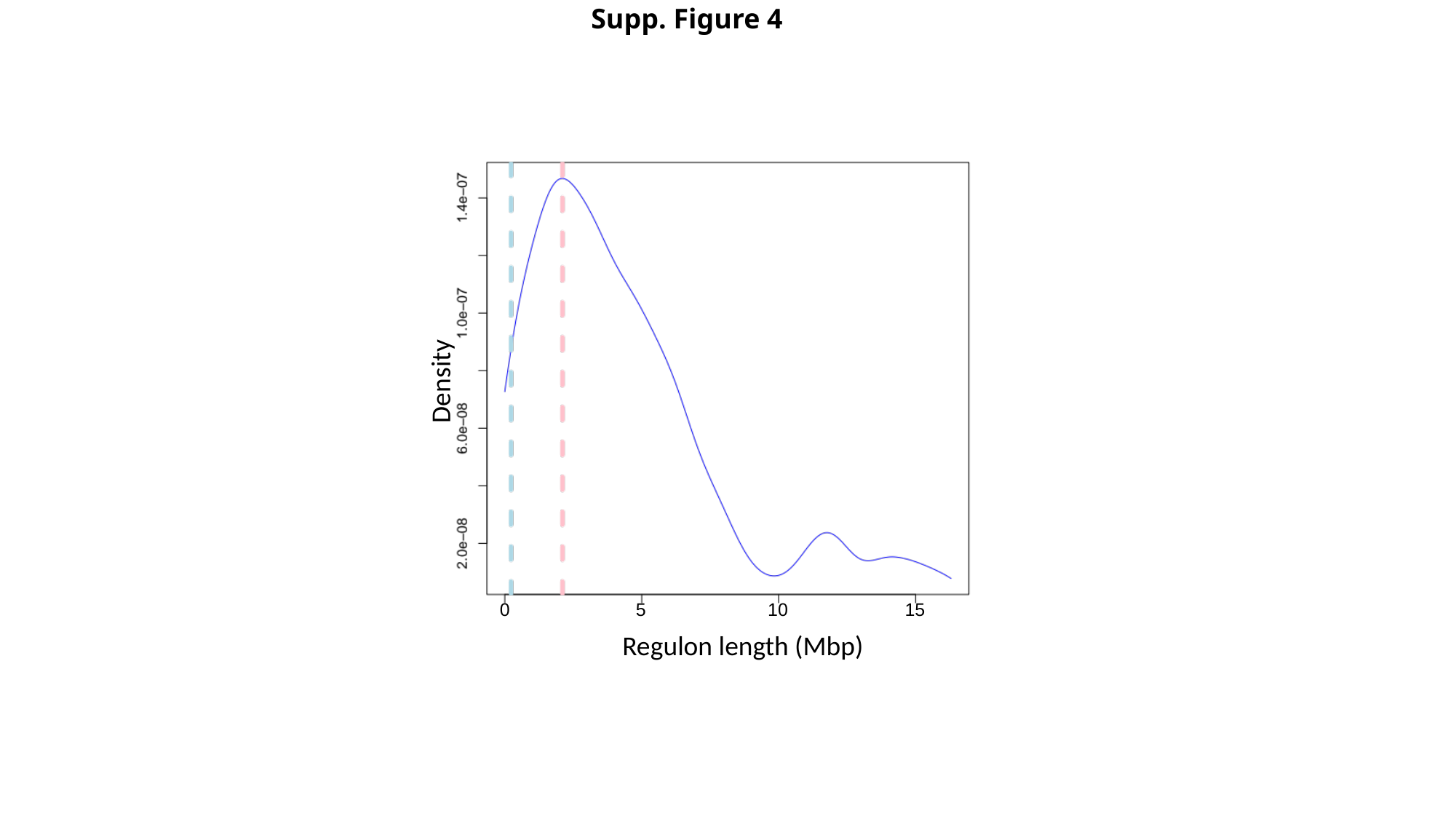

Supp. Figure 4
Density
0
5
10
15
Regulon length (Mbp)

### Slide 13
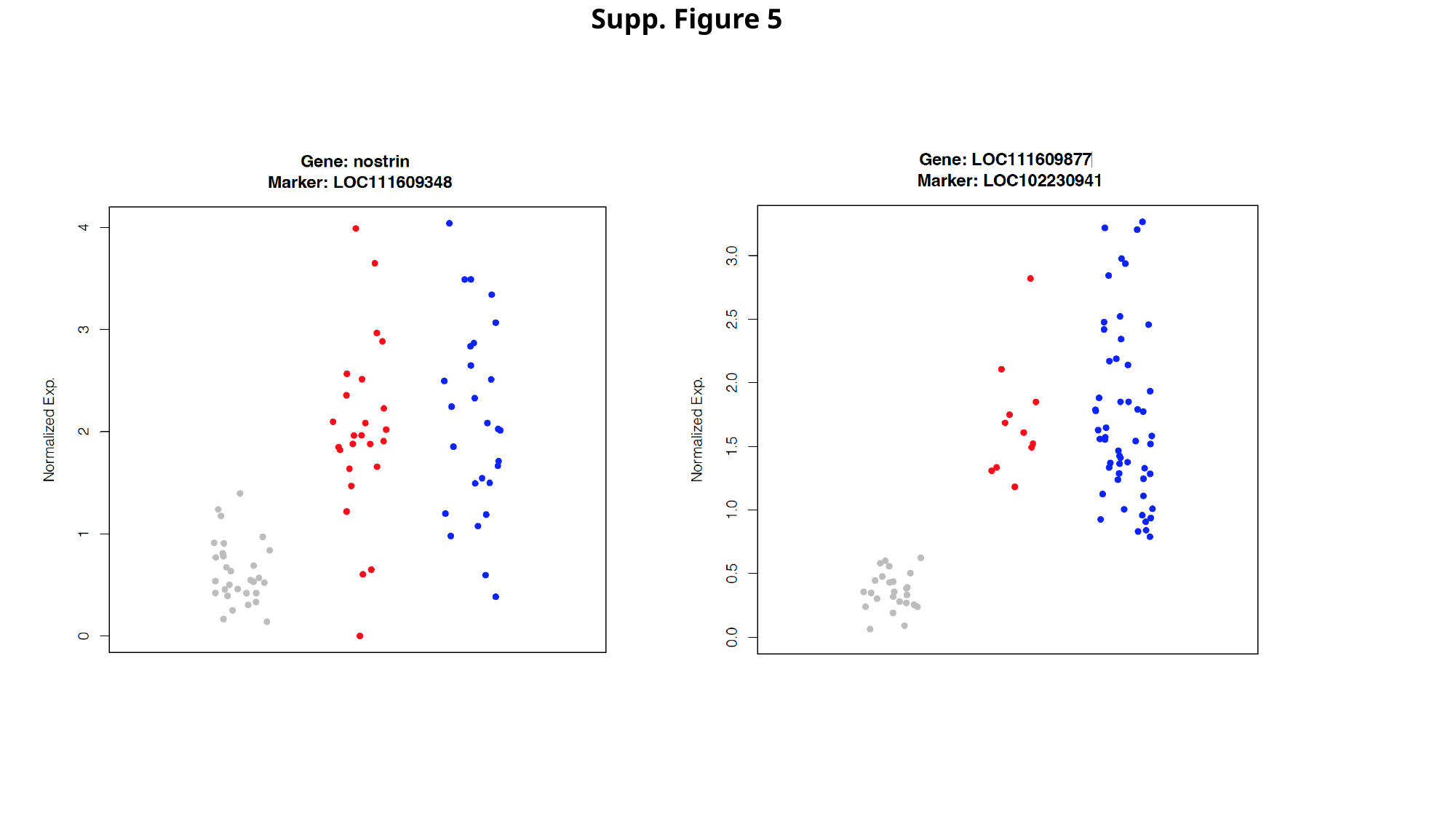

Supp. Figure 5
